# Rsp5/NEDD4 and ESCRT regulate TDP-43 toxicity and turnover via an endolysosomal clearance mechanism

**DOI:** 10.1101/2022.12.05.519172

**Authors:** Aaron Byrd, Lucas Marmorale, Vanessa Addison, Sophia Marcinowski, J. Ross Buchan

## Abstract

A key pathological hallmark in >97% of all Amyotrophic Lateral Sclerosis (ALS) cases is the cytoplasmic mislocalization and aggregation of a nuclear RNA binding protein, TDP-43. Driving clearance of cytoplasmic TDP-43 reduces toxicity in various ALS models, though how TDP-43 clearance is regulated remains controversial. To address this, we conducted an unbiased yeast genome-wide screen using high-throughput dot blots to identify genes that affect TDP-43 levels. Our screen identified ESCRT complex factors, which induce membrane invagination (particularly at multi-vesicular bodies; MVBs) and K63-linked ubiquitination as key facilitators of TDP-43 endolysosomal clearance. TDP-43 co-localized and bound Rsp5/NEDD4 and ESCRT proteins, and perturbations to either increased TDP-43 aggregation and accumulation. NEDD4 also ubiquitinates TDP-43. Lastly, TDP-43 accumulation caused formation of “giant” MVBs which could reflect a pathological consequence of TDP-43 pertinent to ALS. Our studies shed light on endolysosomal-mediated cytoplasmic protein degradation, which likely impacts multiple substrates, and may be a target for novel ALS therapeutic strategies.

## Introduction

Amyotrophic lateral sclerosis (ALS) is an age-related neurodegenerative disease that affects motor neurons and support cells (e.g., glia and astrocytes) of the brain and spinal cord. ALS is characterized by progressive degeneration and death of motor neurons, muscle weakness, and fatal paralysis due to respiratory failure that typically occurs 2-5 years post-diagnosis. While ∼10% of cases exhibit a hereditary component, most ALS cases are sporadic in nature and are not associated with a specific gene mutation. Many familial ALS gene mutations are in genes that impact RNA metabolism and proteostasis, indicating a likely role for these in ALS pathology. However, the molecular mechanisms underlying ALS onset and disease progression are still unknown and no effective treatments for ALS currently exist^1^.

Regardless of disease etiology, a common pathological feature that unites >97% of all ALS cases is the cytoplasmic mislocalization, accumulation, and often aggregation of a nuclear RNA binding protein, TDP-43 (TAR DNA-binding protein 43)^2^. TDP-43 preferentially binds UG-rich sequences, and functions in many RNA-related processes including regulation of alternative splicing, transcription, mRNA localization, translation, mRNA decay, and miRNA biogenesis. TDP-43 mislocalization and accumulation within the cytoplasm results in a toxic gain-of-function, which could reflect TDP-43 aggregates sequestering key RNAs and proteins, as well as a nuclear loss-of-function, in which TDP-43 nuclear depletion may impair the performance of its various nuclear roles^3^.

Failure to effectively clear cytoplasmic TDP-43 may facilitate the pathology of ALS. Indeed, many proteins involved in proteostasis mechanisms such as autophagy and the ubiquitin-proteasome system (UPS), including SQSTM1^4^, VCP^5^, UBQLN2^6^, and OPTN^7^, are mutated in rare familial ALS cases. Also, ALS disease phenotypes can be recapitulated by overexpressing TDP-43 in numerous models (yeast^8^, flies^9^, and human cells^10^) indicating that accumulation of TDP-43 is pathogenic. Additionally, increasing TDP-43 degradation by driving bulk, non-selective autophagy can reverse toxicity associated with TDP-43 accumulation and/or mislocalization^11^. Therefore, increasing clearance of TDP-43 may be a broadly applicable therapeutic strategy for diseases with cytoplasmic TDP-43 proteinopathy.

The mechanisms by which TDP-43 is degraded remain highly debated. Prior studies have postulated that insoluble TDP-43 aggregates and oligomers are targeted for clearance via autophagy and soluble TDP-43 is degraded via the UPS^12–14^. However, these studies often relied upon chemical means of blocking autophagy (Bafilomycin A1, Wortmannin, 3-methyladenine, Chloroquine, etc.) which also impair other lysosomal trafficking pathways (e.g., endolysosomal trafficking), or lysosome function generally, making the interpretation of results difficult. Several E3-ubiquitin ligases can ubiquitinate TDP-43, but the effects of these ubiquitination events are often not correlated to TDP-43 clearance^15–18^.

We and others have shown in yeast, human cell culture, and iPSC ALS patient derived motor neurons that cytoplasmic TDP-43 clearance and toxicity are strongly dependent on an autophagy-independent endolysosomal degradation pathway ^19–22^. Notably, blocks to autophagy and the proteasome showed minimal effects on TDP-43 clearance and toxicity in yeast models. In yeast and human cell models, impairment of endolysosomal flux caused increased TDP-43 accumulation, aggregation, and toxicity, whereas enhancing endolysosomal flux suppressed these phenotypes. In a TDP-43 fly model of ALS, genetic defects in endocytosis exacerbated motor neuron dysfunction whereas enhancement of endocytosis suppressed such dysfunction. Lastly, expression of cytoplasmic TDP-43, in particular aggregation prone mutants, inhibits endocytosis^19, 20^.

Given the lack of consensus on how TDP-43 is cleared, and which clearance pathways are most relevant to ALS phenotypes, we performed an unbiased genetic screen in *S. cerevisiae* to determine genes that alter TDP-43 abundance. We discovered additional conserved proteins that regulate TDP-43 toxicity, aggregation, and protein stability by targeting TDP-43 for endolysosomal clearance, specifically Endosomal Sorting Complexes Required for Transport (ESCRT) proteins, and the E3 ubiquitin ligase, Rsp5 (yeast)/NEDD4 (human). NEDD4 also binds, ubiquitinates and destabilizes TDP-43. Finally, mild TDP-43 over-expression (2-fold including endogenous TDP-43) causes accumulation of giant MVB-like organelles, within which TDP-43 accumulates in a NEDD4-driven manner. Collectively, these data and prior work suggest that cytoplasmic TDP-43 is cleared by an endolysosomal mechanism, and that pathological accumulation of TDP-43 can also disrupt endolysosomal trafficking.

## Results

### A dot-blot genetic screen in yeast identifies genes regulating TDP-43 steady-state levels

To our knowledge, an unbiased genetic screen to identify regulators of TDP-43 abundance has not been conducted in any model system. Such a screen could identify regulators of both TDP-43 synthesis and degradation, and in turn novel therapeutic targets for diseases characterized by TDP-43 proteinopathy. Therefore, we designed a dot blot genetic screen, a refinement of a previously published method^23^, that allowed us to measure steady-state TDP-43 protein levels in each strain of the yeast non-essential gene deletion library (**Figure 1A;** see *Materials and Methods*). To control for variations in expression due to intrinsic or extrinsic noise, four control strains were utilized in two different locations on each 384-well dot blot plate. First, a wild-type (WT) strain containing TDP-43-YFP plasmid was included to act as the standard to which all other strains were normalized. Second, a *bre5Δ* strain containing TDP-43-YFP plasmid was used to ensure that strains with lower TDP-43 protein levels were detectable.

**Figure 1:**
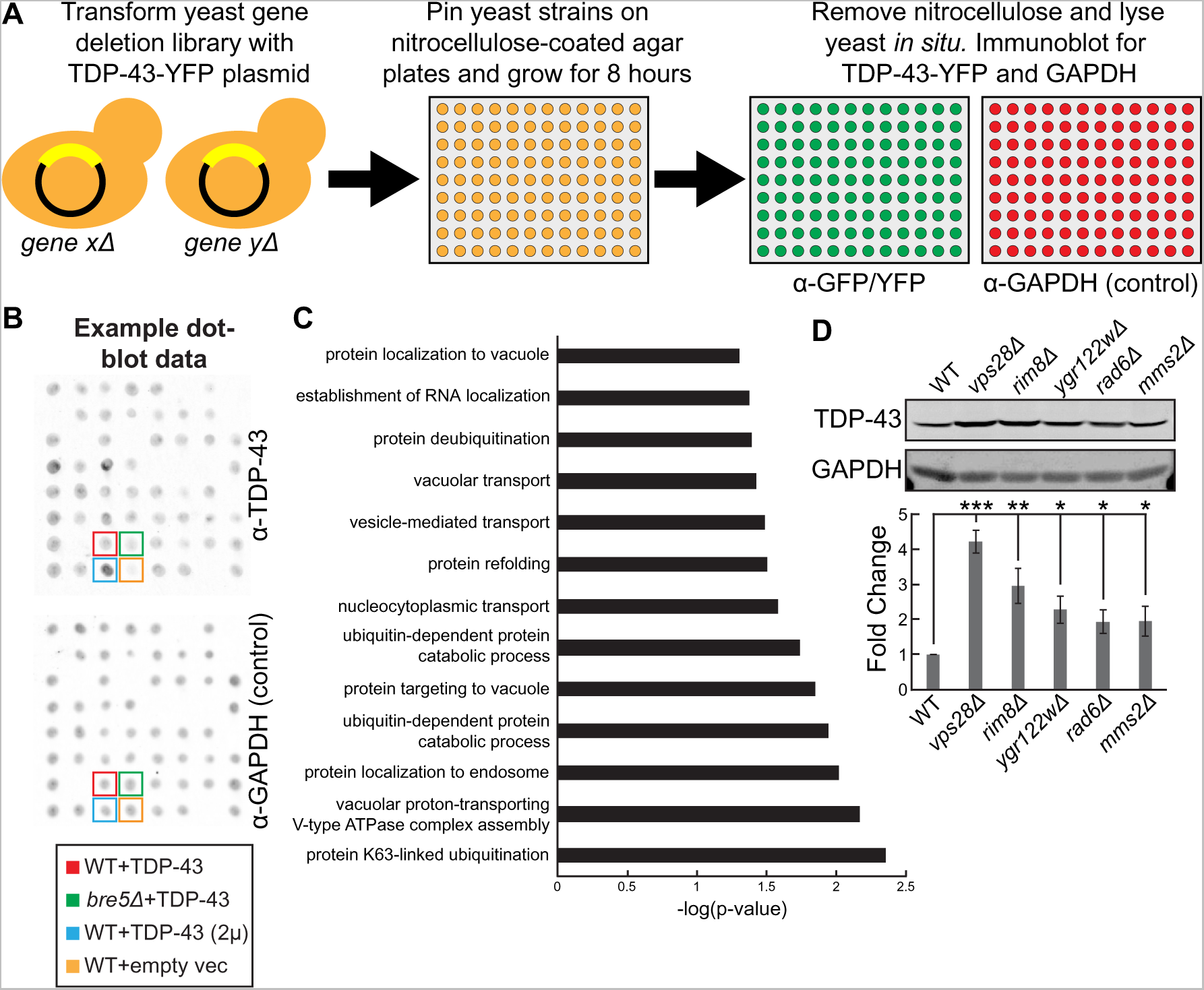
A dot-blot genetic screen in yeast identifies genes regulating TDP-43 steady-state levels. (A) Schematic representation of the dot-blot genetic screen. (B) Example dot-blot with control strains and placement indicated. (C) Gene ontology terms of biological functions significantly enriched within the hits from the dot-blot genetic screen. (D) Indicated strains were transformed with TDP-43-mRuby2 plasmid and grown to mid-logarithmic growth phase and TDP-43-mRuby2 was detected by western blot using anti-TDP-43 antibody. TDP-43 levels were normalized to GAPDH loading control. *, *P* < 0.05; **, *P* < 0.01; ***, *P* < 0.001 by analysis of variance with Tukey’s *post hoc* test; *N* = 3

We previously reported that the *bre5Δ* strain (a deubiquitinase co-factor) exhibits significantly reduced TDP-43 protein levels at steady-state^20^. Third, a WT strain containing a high-copy TDP-43-GFP plasmid was also included to ensure strains with higher TDP-43 protein levels were detectable. Lastly, a WT strain containing an empty vector was included to ensure that nonspecific binding of the α-GFP/YFP antibody was minimal **(Figure 1B)**. Besides α-GFP/YFP labeling, blots were also probed with an α-GAPDH antibody, which allowed normalization of each dot blot and mitigated variable detection caused by variable strain growth rates or lysis efficiencies **(Figure 1B)**.

Altogether, this screen generated 196 hits total with 44 strains exhibiting significantly increased TDP-43 abundance (>1.5-fold increase, p-value < 0.05) and 152 strains exhibiting significantly decreased TDP-43 abundance (<0.5 fold decrease, p-value <0.05; **Supplementary Table 1**). Encouragingly, our screen re-identified previously known regulators of TDP-43 abundance in yeast such as *bre5Δ* (30% of WT)^20^. Other expected regulators of TDP-43 abundance obtained in the screen included regulators of carbon metabolism, consistent with the *GAL1-*driven nature of the TDP-43 construct utilized. Furthermore, two separate *rim8Δ* strains, present twice in the deletion library owing to a gene annotation error, were both identified as hits with very similar levels of TDP-43 accumulation (5.0- and 5.5-fold accumulation).

Overall, gene ontology analysis indicated that terms most significantly enriched among the hits from the screen included K63-linked ubiquitination, vacuolar proton-transporting V-type ATPase complex assembly, and protein localization to endosomes **(Figure 1C)**. Interestingly, among the top 10 hits with increased TDP-43 abundance were *rim8Δ*, *mms2Δ*, *vps28Δ*, *ygr122wΔ*, and *rad6Δ* strains, which we validated manually via western blotting **(Figure 1D)**. Rim8 is an adaptor in the endolysosomal pathway for the E3 ligase Rsp5, which facilitates targeting of Rsp5 to its substrates for ubiquitination^24^. Rim8 also binds the ESCRT-I complex to target its substrates (typically transmembrane proteins) for eventual degradation in the vacuole^25^. Mms2 and Rad6 are E2 ubiquitin-conjugating enzymes that both function in K63-linked protein ubiquitination^26, 27^. Vps28 is a core component of the ESCRT-I complex involved in sorting cargoes into MVBs^28^. Ygr122w is a protein of unclear function that binds the ESCRT-III complex factor Snf7, and negatively genetically interacts with Rsp5^29^. Together, combined with our prior work^19, 20^, this led us to a working model that TDP-43 is subject to K63-linked ubiquitination via Rsp5, which then ultimately targets it to the ESCRT complex for internalization into MVBs.

### TDP-43 clearance and aggregation depend on ESCRT function

Previously, we showed that mutations in ESCRT genes increase TDP-43 toxicity^19^. To verify the defects in TDP-43 clearance in ESCRT mutants as suggested by our screen, we measured protein levels of TDP-43 in mutants of each ESCRT subcomplex via western blotting and found that almost all these mutant strains had significantly increased steady-state levels of TDP-43 when compared to a wild-type strain (**Figure 2A**). We then sought to determine if these increased levels of total TDP-43 protein correlated with an increase in TDP-43 aggregation via fluorescence microscopy. Indeed, relative to WT, strains that exhibited small increases in TDP-43 abundance also had small increases in TDP-43 foci per cell (*vps4Δ* and *vps2Δ*), while strains that had large increases in TDP-43 abundance had significantly increased TDP-43 foci per cell (*vps28Δ*, *vps36Δ*, and *vps27Δ*; **Figure 2B)**. Additionally, to determine whether increases in TDP-43 abundance were a result of improper clearance, rather than altered synthesis, we performed cycloheximide chase assays where global protein synthesis was halted by the addition of cycloheximide and then yeast whole cell lysate was collected at different timepoints over 24 hours to determine how quickly TDP-43 is cleared. In these assays, TDP-43 stability was significantly increased in the ESCRT mutant strains *vps36Δ* and *vps4Δ* **(Figure 2C)**. TDP-43 foci also colocalized with the core ESCRT factors Vps28, Vps36, and Vps4 indicating a potential physical interaction between TDP-43 and the ESCRT complex **(Figure 2D)**. To assess this, we performed immunoprecipitations of GFP-tagged ESCRT factors Vps28 (ESCRT-I) and Vps36 (ESCRT-II) to determine if TDP-43 can interact with these proteins. We found that TDP-43 co-immunoprecipitated with both Vps28 and Vps36 indicating that TDP-43 can indeed interact with the ESCRT complex in yeast cells **(Figure 2E)**. Altogether, these data strongly suggest that TDP-43 clearance in yeast is dependent upon the ESCRT complex.

**Figure 2:**
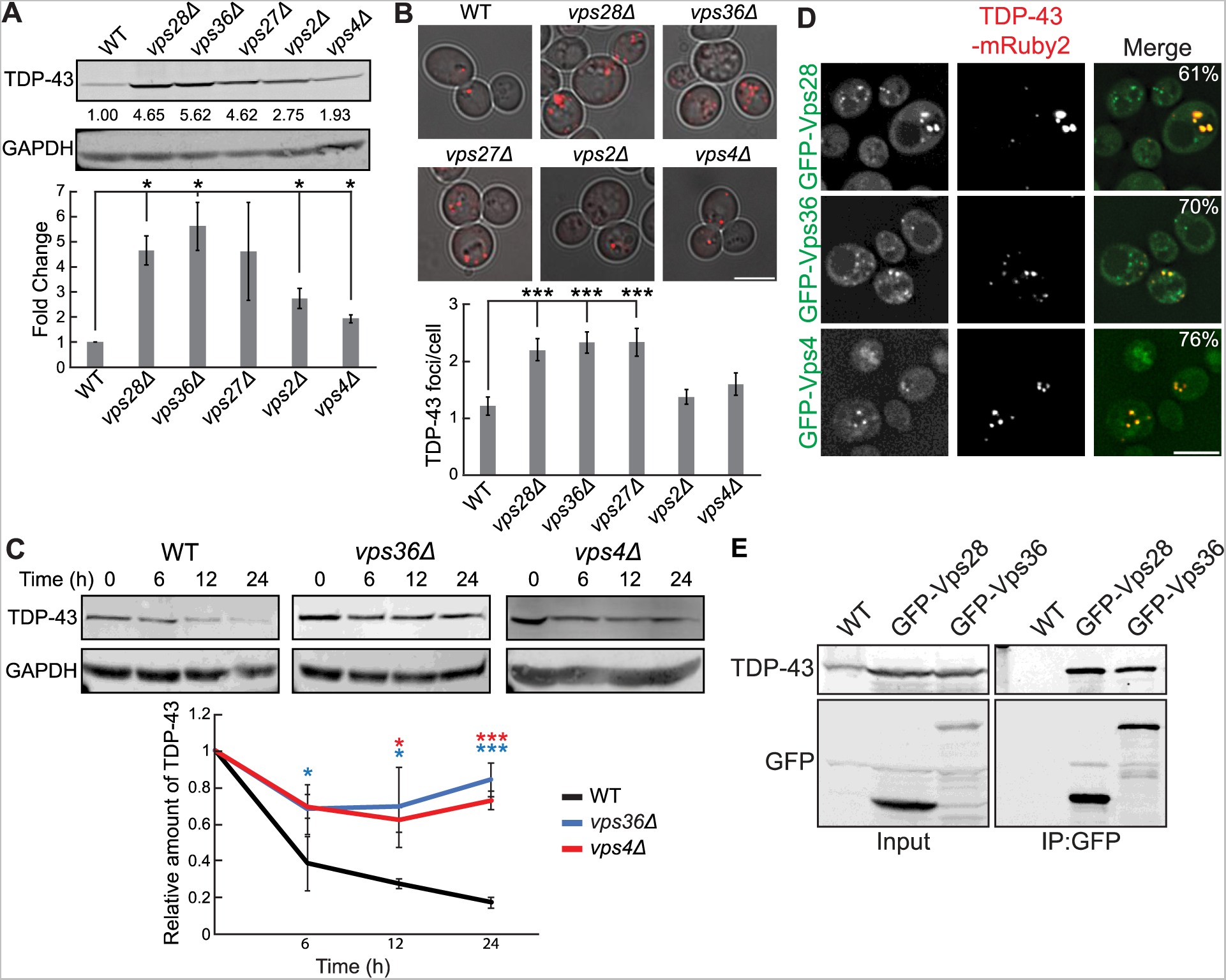
TDP-43 clearance and aggregation depend on ESCRT function. (A) Indicated strains were transformed with TDP-43-mRuby2 plasmid and grown to mid-logarithmic growth phase and TDP-43-mRuby2 was detected by western blot using anti-TDP-43 antibody. TDP-43 levels were normalized to GAPDH loading control. *, *P* < 0.05 by analysis of variance with Tukey’s *post hoc* test; *N* = 3 (B) Indicated strains were transformed with TDP-43-mRuby2 plasmid and imaged in mid-logarithmic growth phase with foci/cell quantification below. *P* < 0.001 analysis of variance with Tukey’s *post hoc* test; *N* = 3; Bar, 5 µm. (C) Indicated strains were transformed with TDP-43-mRuby2 plasmid and grown to mid-logarithmic growth phase before being treated with 0.2 mg/ml cycloheximide for 0, 6, 12, and 24 hours. TDP-43-mRuby2 levels were detected by western blot using anti-TDP-43 antibody and normalized to GAPDH loading control. *, P < 0.05; ***, *P* < 0.001 analysis of variance with Tukey’s *post hoc* test; *N* = 3. (D) Strains endogenously expressing GFP-Vps28, GFP-Vps36, and GFP-Vps4 were transformed with a TDP-43-mRuby2 plasmid and imaged at mid-log phase growth. Percentage values indicate colocalization frequency of TDP-43-mRuby2 foci with GFP foci. *N* = 3; Bar, 5 µm (E) Strains with endogenously expressed GFP-Vps28 or GFP-Vps36 and the isogenic WT strain were transformed with TDP-43-mRuby2 and grown to mid-logarithmic growth phase before GFP immunoprecipitation was performed. *N* = 3.

### TDP-43 toxicity, aggregation, and clearance depend on the E3-ubiquitin ligase Rsp5

Since factors related to K63 ubiquitination and known Rsp5 interactors were among the top hits of our dot blot screen, we decided to investigate the role of the E3 ubiquitin-ligase Rsp5 in the toxicity, aggregation, and clearance of TDP-43. Rsp5 is one of the major endolysosomal E3 ubiquitin-ligases in yeast responsible for K63-linked ubiquitination and has many functions in regulating protein turnover and trafficking^30^. Importantly, *RSP5* is an essential gene, thus we utilized a commonly used temperature-sensitive (*ts*) mutant, *rsp5-3*, which possesses three missense point mutations, T104A, E673G, and Q716P, that inhibit Rsp5’s catalytic activity at elevated temperatures^31^. TDP-43 toxicity, evidenced via serial dilution spotting assays, was greatly exacerbated by partial inactivation of *rsp5-3* by temperature shift to 30°C **(Figure 3A)**. Surprisingly, even at the “permissive” temperature of 25°C, the *rsp5-3* strain exhibited greater growth defects from TDP-43 expression, which may reflect partially defective Rsp5 function. Next, we examined if enhanced TDP-43 toxicity in the *rsp5-3* strain correlated with an increase in TDP-43 accumulation and aggregation. As expected, we saw an increase in TDP-43 aggregation when *rsp5-3* was strongly inactivated by shifting the temperature to 35°C while comparable amounts were seen at the permissive temperature of 25°C **(Figure 3B)**. We next sought to determine whether Rsp5 function is important for the clearance of TDP-43 by performing cycloheximide chase assays. In the wild-type and *rsp5-3* strains grown at 25°C, TDP-43 abundance decreased similarly over a 24hr time course. However, when strains were shifted to 35°C, the *rsp5-3* strain exhibited significantly greater TDP-43 stability than the wild-type strain **(Figure 3C)**. Additionally, Rsp5 co-localizes in TDP-43 cytoplasmic foci **(Figure 3D)** and a physical interaction between both proteins was detected via immunoprecipitation **(Figure 3E)**. Notably, when we overexpressed Rsp5 in a WT strain, TDP-43 toxicity and steady-state protein levels were significantly reduced **(Figures 3F, 3G)**.

**Figure 3:**
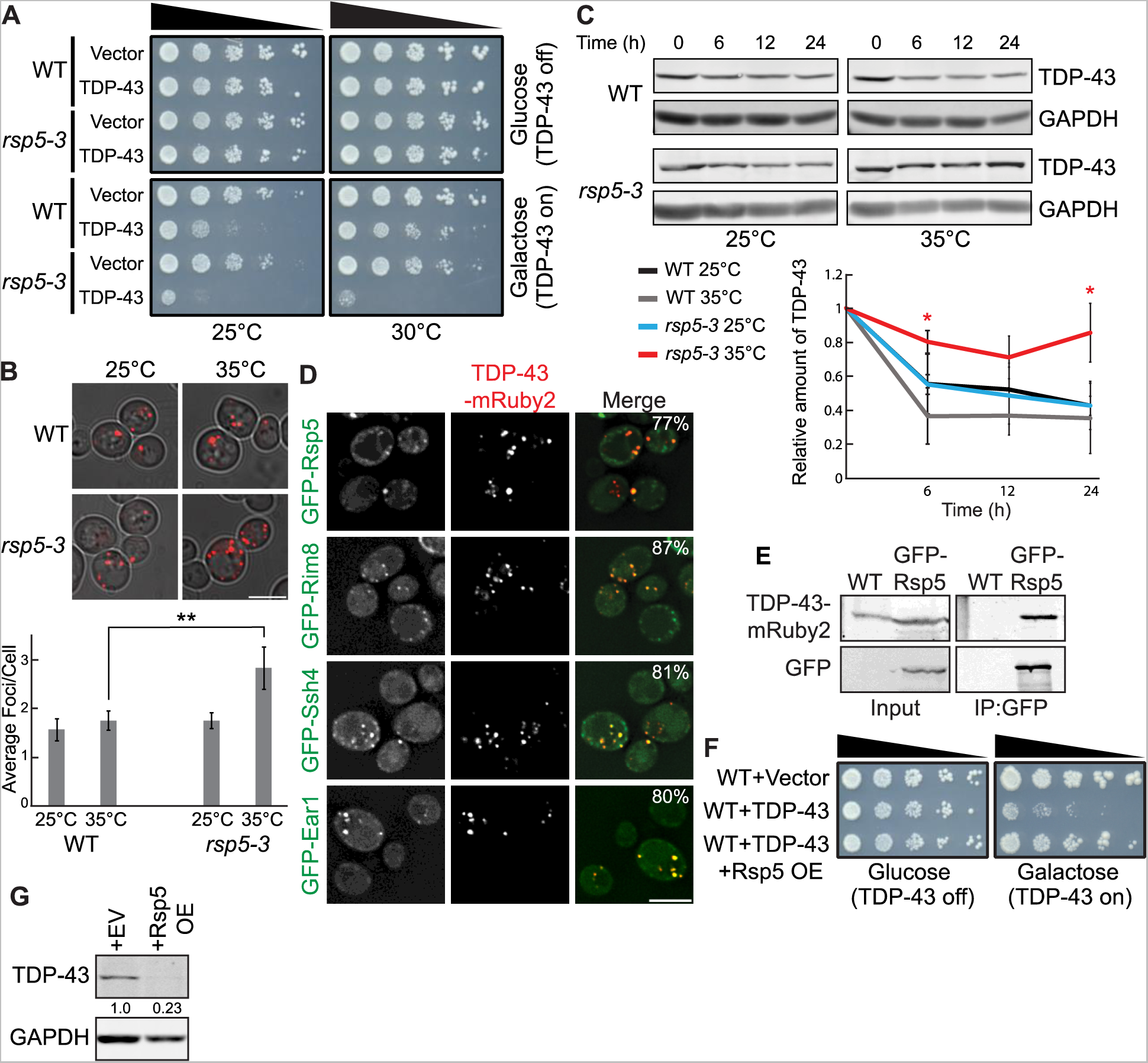
TDP-43 toxicity, aggregation, and toxicity depend on the E3 ligase Rsp5. (A) Representative serial dilution growth assay of WT and *rsp5-3* strains transformed with the empty vector or untagged TDP-43 plasmid grown at 25°C or 30°C as indicated. *N* = 3 (B) WT and *rsp5-3* strains were transformed with TDP-43-mRuby2 plasmid and grown to mid-logarithmic growth phase at 25°C and samples were imaged. Cultures were then incubated at 35°C for 16 hours and imaged again. **, *P* < 0.01 by analysis of variance with Tukey’s *post hoc* test; *N* = 3; Bar, 5 µm (C) WT and *rsp5-3* strains were transformed with TDP-43-mRuby2 plasmid and grown to mid-logarithmic growth phase at 25°C before being treated with 0.2 mg/ml cycloheximide for 0, 6, 12, and 24 hours. At time 0 cultures were either shifted to 35°C or kept at 25°C for the remainder of the time course. TDP-43-mRuby2 levels were detected by western blot using anti-TDP-43 antibody and normalized to GAPDH loading control. *, *P* < 0.05 by analysis of variance with Tukey’s *post hoc* test; *N* = 3 (D) Strains endogenously expressing GFP-Rsp5, GFP-Rim8, GFP-Ssh4 and GFP-Ear1 were transformed with a TDP-43-mRuby2 plasmid and imaged at mid-log phase growth. Percentage values indicate colocalization frequency of TDP-43-mRuby2 foci with GFP foci. *N* = 3; Bar, 5 µm (E) A strain endogenously expressing GFP-Rsp5 and the isogenic WT strain were transformed with TDP-43-mRuby2 and grown to mid-logarithmic growth phase before GFP immunoprecipitation was performed. *N* = 3 (F) Representative serial dilution growth assay of WT strain transformed with an empty vector control or untagged TDP-43 plasmid, and with the empty vector control or Rsp5 overexpression plasmid. *N* = 3 (G) WT strain was transformed with TDP-43-mRuby2 and then also transformed with Rsp5 overexpression or empty vector control plasmid and grown to mid-logarithmic growth phase. TDP-43-mRuby2 was detected by western blot using anti-TDP-43 antibody. TDP-43 levels were normalized to GAPDH loading control. *, *P* < 0.05 by analysis of variance with Tukey’s *post hoc* test. *N* = 3

Rsp5 substrate specificity often requires adaptor proteins of the arrestin-related trafficking adaptor family^30^. Among the best-known Rsp5 adaptors are Rim8, Ear1, and Ssh4 which are also all involved in helping sort Rsp5 substrates to the endolysosomal pathway^30^. Rim8 was a top repeated hit in our dot blot screen, having a >5-fold increase in TDP-43 abundance (**Supplementary Table 1**). Ear1 and Ssh4 often function redundantly but both are involved in cargo sorting of proteins delivered to MVBs and were therefore of interest to our study. TDP-43 colocalizes with all three of these Rsp5 adaptor proteins via fluorescence microscopy (**Figure 3D**). To comprehensively address the importance of all Rsp5 adaptors in regulation of TDP-43 levels, we performed steady state western blotting of TDP-43 abundance in all 21 Rsp5 adaptor nulls. Only *rim8Δ* showed a significant effect on TDP-43 clearance (**Figure S1**). In summary, Rsp5 regulates TDP-43 turnover and toxicity in our yeast model, aided in part by the adaptor Rim8.

### Dominant negative VPS4^E228Q^ increases TDP-43 stability and promotes cytoplasmic mislocalization

To examine the role of ESCRT-dependent turnover of TDP-43 in human cells, we utilized HEK293A cells and a dominant negative mutant of the core ESCRT protein, VPS4. This mutant possesses a point mutation, E228Q, that prevents ATP-hydrolysis and greatly hinders ESCRT functions^32^. We first determined whether EGFP-VPS4^E228Q^ expression affected the turnover of endogenous TDP-43 by performing cycloheximide chase assays. We found that endogenous TDP-43 in cells transfected with EGFP-VPS4^E228Q^ was significantly more stable than untreated WT cells **(Figure 4A)**. Since TDP-43 colocalized with VPS4 in yeast, we investigated whether TDP-43 colocalized with EGFP-VPS4^E228Q^ by performing immunofluorescence of endogenous TDP-43. We observed strong colocalization of TDP-43 with EGFP-VPS4^E228Q^ in large cytoplasmic foci some of which took on vesicular-like patterns of colocalization **(Figure 4B; see middle row)**. Endogenous TDP-43 levels in both nuclei and the cytoplasm also increased significantly in the VPS4 ^E228Q^ background (**Figure 4B-C**) TDP-43 was significantly more likely to form cytoplasmic foci in cells transfected with EGFP-VPS4^E228Q^ versus WT cells **(Figure 4D)**. Lastly, we observed that the decreased cell viability associated with overexpression of TDP-43-GFP and or the C-terminal fragment (CTF) and disease-associated isoform, TDP-35-GFP, was significantly exacerbated by expression of VPS4^E228Q^ **(Figure 4E)**. Collectively, these data suggest that the ESCRT complex facilitates TDP-43 turnover and in turn regulates TDP-43 toxicity.

**Figure 4:**
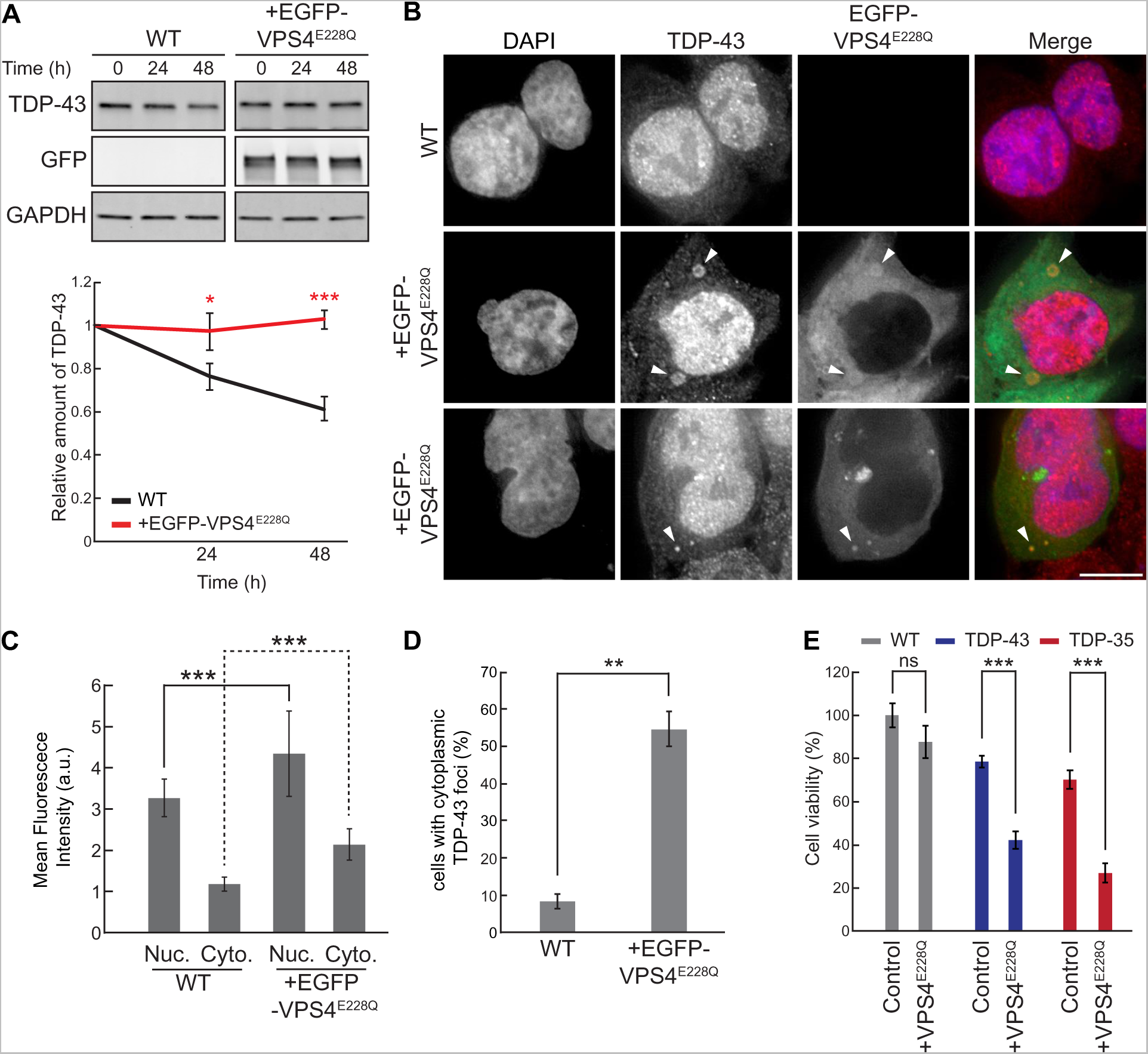
VPS4^E228Q^ increases TDP-43 stability and promotes cytoplasmic mislocalization. (A) HEK293A WT cells were transfected with EGFP-VPS4^E228Q^ plasmid or empty vector control and grown in normal conditions for 48 hours. Cells were then treated with 0.3 mg/ml cycloheximide for the indicated amounts of time and TDP-43 levels were detected by western blot with anti-TDP-43 antibodies and normalized to the level of the GAPDH loading control. *, *P* < 0.05; ***, *P* < 0.001 by analysis of variance with Tukey’s *post hoc* test; *N* = 3 (B) HEK293A WT cells were transfected with empty vector control (top row) or EGFP-VPS4^E228Q^ plasmid (next two rows, to show phenotypic diversity) and grown in normal conditions for 48 hours before being fixed and immunostained with anti-TDP-43 antibodies. *N* = 3; Bar, 10 µm (C) Quantification of mean nuclear and cytoplasmic TDP-43 signal intensities in HEK293A WT cells and cells transfected with EGFP-VPS4^E228Q^ plasmid. ***, *P* < 0.001 by analysis of variance with Tukey’s *post hoc* test; *N* = 30 (D) Quantification of the percentage of cells with cytoplasmic TDP-43 foci in HEK293A WT cells and cells transfected with EGFP-VPS4^E228Q^ plasmid. **, *P* < 0.01 by student’s two-tailed T-test; *N* = 3 (E) HEK293A, HEK-TDP-43-GFP, and HEK-TDP-35-GFP cells were seeded at equal cell numbers and transfected with VPS4^E228Q^ plasmid or empty vector control and cell viability was compared after 48 hours. ***, *P* < 0.001 by analysis of variance with Tukey’s *post hoc* test; *N* = 3

### NEDD4 regulates TDP-43 stability and ubiquitination

We next determined if NEDD4, the Rsp5 homolog in humans, affects clearance of TDP-43 in a mild overexpression context. We utilized HEK293A cells stably expressing TDP-43-GFP at comparable levels to endogenous TDP-43 in HEK293A cells; this also results in a slight increase in cytoplasmic TDP-43 localization^19^. By performing a cycloheximide chase assay, we found that TDP-43-GFP stability was significantly increased in cells where NEDD4 expression was knocked down approximately 50-60% (depending on the timepoint) versus WT **(Figure 5A)**. Additionally, overexpression of NEDD4 via NEDD4-mCherry plasmid transfection significantly lowered the steady-state levels of TDP-43-GFP **(Figure 5B)**. Taken together, these data indicate that TDP-43 stability and abundance are regulated by NEDD4. TDP-43 cytoplasmic abundance tends to correlate with induced cellular toxicity, therefore we investigated whether altering TDP-43 levels and stability by overexpression or knockdown of NEDD4 affected cell viability. We found that the decreased cell viability associated with overexpression of TDP-43-GFP and TDP-35-GFP was significantly rescued by overexpression of NEDD4 **(Figure 5C)**. When NEDD4 was knocked down we found that TDP-43-GFP and TDP-35-GFP expression significantly reduced cell viability **(Figure 5C)**. Thus, NEDD4 is a regulator of TDP-43 toxicity.

**Figure 5:**
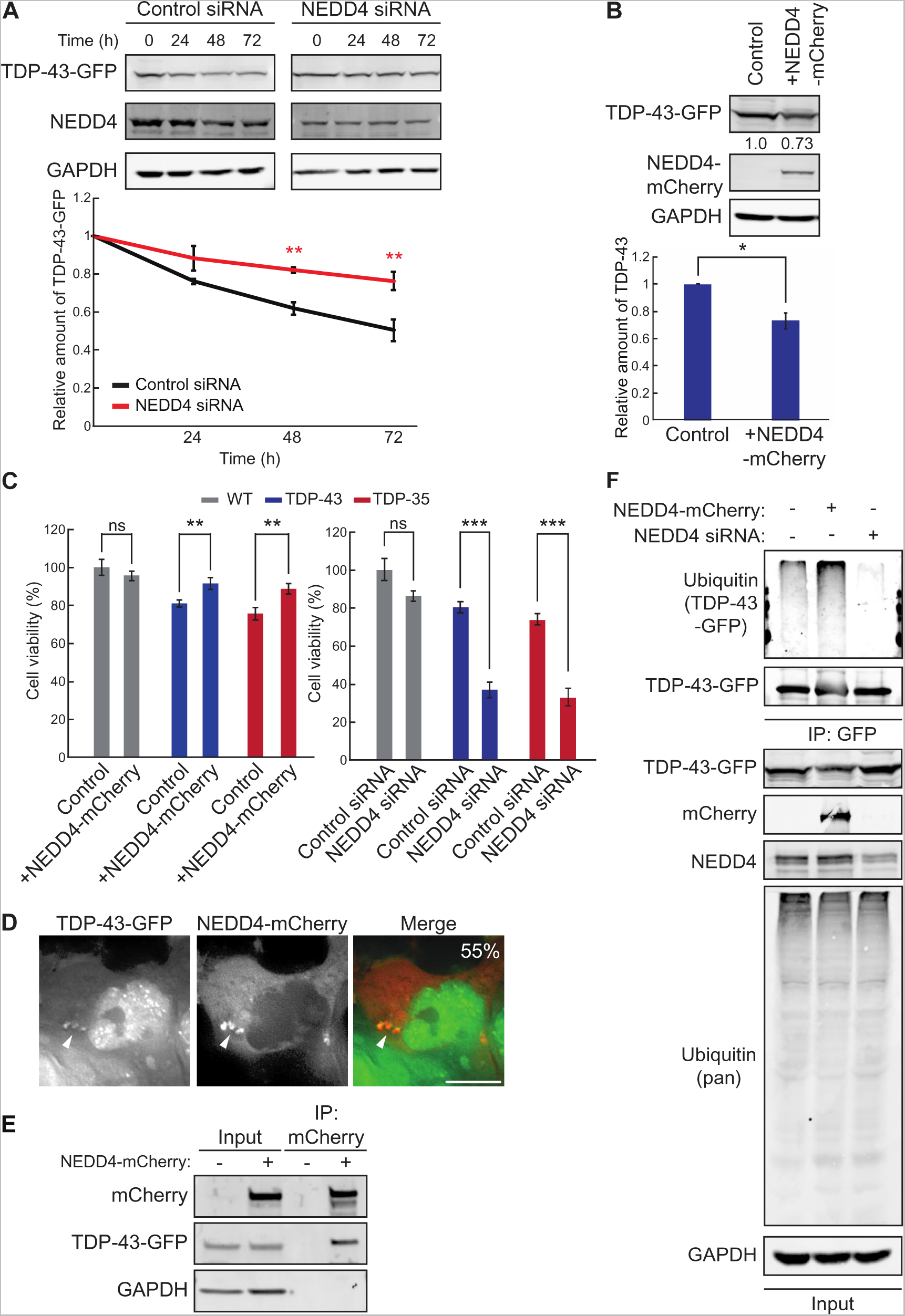
NEDD4 regulates TDP-43 turnover, toxicity, and ubiquitination. (A) HEK-TDP-43-GFP c ells vvere transf e cted vvtth stRNAs targ e ting NEDD4 or non-codtng c ontrol and grown in normal conditions for 3 days. Cells were then treated with 0.3 mg/ml cycloheximide for the indicated amounts of time and TDP-43-GFP levels were detected by western blot with anti-GFP antibodies and normalized to the level of the GAPDH loading control. **, *P* < 0.01 by analysis of variance with Tukey’s *post hoc* test; *N* = 3 (B) HEK-TDP-43-GFP cells were transfected with NEDD4-mCherry plasmid and grown in normal conditions for 3 days. TDP-43-GFP levels were detected by western blot with anti-GFP antibodies and normalized to the level of the GAPDH loading control. *, *P* < 0.05; by student’s two-tailed T-test; *N* = 3. (C) HEK293A, HEK-TDP-43-GFP, and HEK-TDP-35-GFP cells were seeded at equal cell numbers and transfected with NEDD4-mCherry or empty vector control (left), or siRNA targeting NEDD4 or non-coding control (right) and cell viability was compared after 48 hours. **, *P* < 0.01; ***, *P* < 0.001 by analysis of variance with Tukey’s *post hoc* test; *N* = 3 (D) HEK-TDP-43-GFP cells were transfected with NEDD4-mCherry plasmid and incubated for 24 hours before being imaged. Percentage value indicates colocalization frequency of cytoplasmic TDP-43-GFP foci with NEDD4-mCherry foci. *N* = 3; Bar, 10 µm (E) HEK-TDP-43-GFP cells were transfected with NEDD4-mCherry plasmid or empty vector and incubated for 48 hours before mCherry was immunoprecipitated and TDP-43-GFP binding was assessed via western blot using anti-TDP-43 antibodies. (F) HEK-TDP-43-GFP cells were transfected with siRNAs targeting NEDD4, NEDD4-mCherry plasmid, or a mock control and grown for 3 days before TDP-43-GFP was immunoprecipitated under denaturing conditions (See *Materials and Methods*).

To determine whether the effects of NEDD4 overexpression involved interaction with TDP-43, we initially performed live-cell fluorescence microscopy of cells overexpressing NEDD4-mCherry and TDP-43-GFP. We found that NEDD4-mCherry and TDP-43-GFP colocalize in cytoplasmic foci **(Figure 5D)**. *In vivo* binding of NEDD4-mCherry and TDP-43-GFP was also detected via immunoprecipitation **(Figure 5E)**. Finally, since NEDD4 is an E3 ligase, and ubiquitination is commonly involved in protein degradation, we performed a ubiquitination assessment of immunopurified TDP-43-GFP from cells with NEDD4 overexpression or NEDD4 knockdown. In NEDD4 overexpression cells, we observed an increase in the ubiquitination of TDP-43-GFP versus control **(Figure 5F)**. Conversely, in NEDD4 siRNA knockdown cells, TDP-43-GFP ubiquitination was markedly decreased **(Figure**

### TDP-43 accumulation causes enlargement of MVBs

Since our working model of TDP-43 clearance via the ESCRT machinery suggests TDP-43 localizes within MVBs, we decided to investigate this via immunofluorescence microscopy using the common MVB marker, Rab7. Surprisingly, we discovered that TDP-43-GFP and TDP-35-GFP cell lines developed “giant MVBs” **(Figure 6A)**. Where typically MVBs reach about 0.5-1μm in diameter^33^, we commonly observed Rab7 structures that were much larger in diameter. These giant MVBs, defined as Rab7 structures with diameters >2.5μm, were significantly enriched in TDP-43-GFP and TDP-35-GFP cells versus parental HEK293A cells **(Figure 6B)** and averaged 4.5-5μm in diameter **(Figure 6C)**. Rab7-positive MVBs in our TDP-43-GFP expressing cell lines are highly variable in size with some reaching as large as >10μm in diameter **(Figure S2A)**.

**Figure 6:**
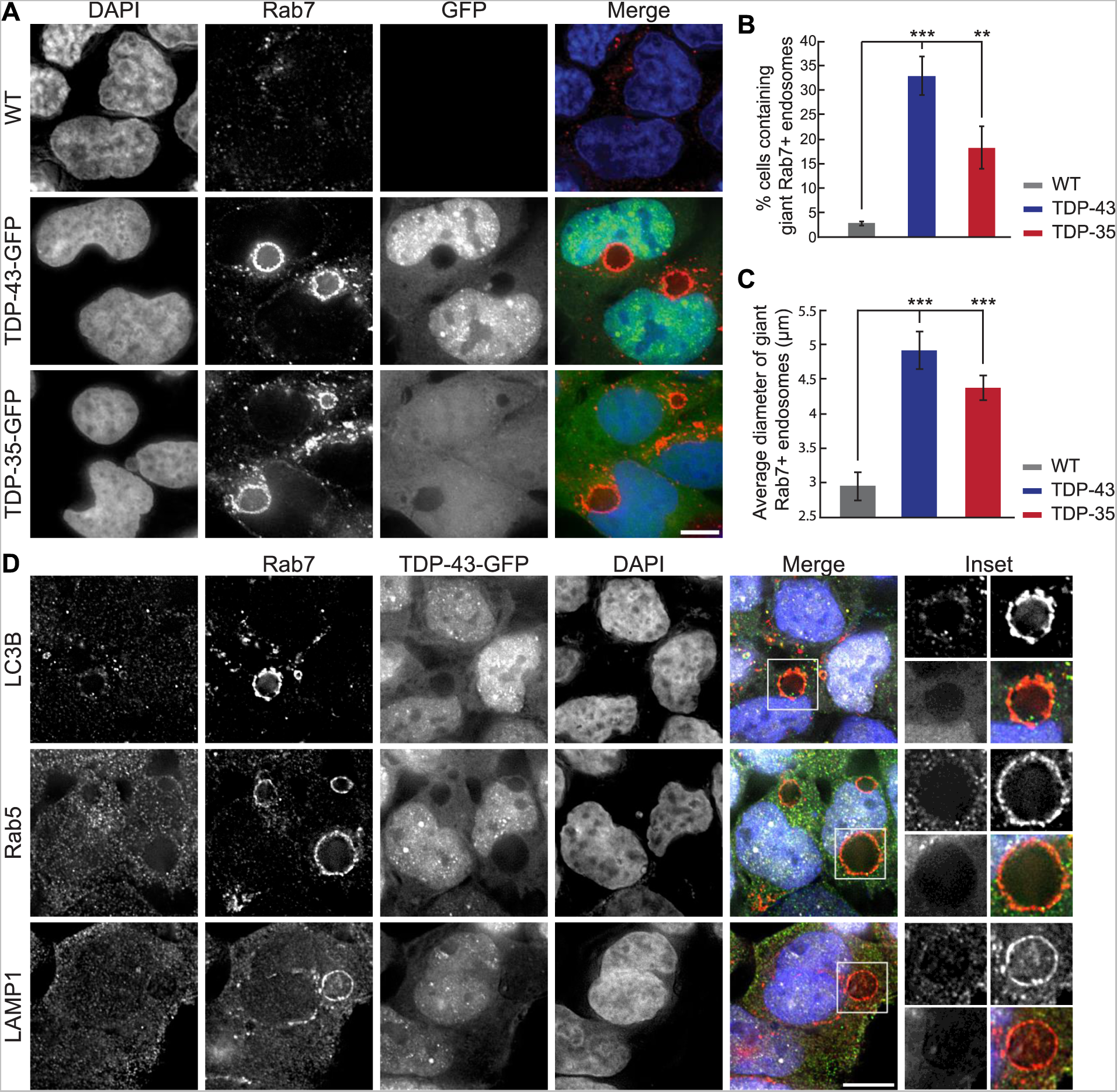
TDP-43 accumulation causes enlargement of MVBs. (A) HEK293A WT, HEK-TDP-43-GFP, and HEK-TDP-35-GFP cells were grown in normal conditions before being fixed and immunostained for Rab7. *N* = 3; Bar, 10 μm (B) Quantification of the frequency of giant endosomes in HEK293A WT, HEK-TDP-43-GFP, and HEK-TDP-35-GFP cells. **, *P* < 0.01; ***, *P* < 0.001 by analysis of variance with Tukey’s *post hoc* test; *N* = 3 (C) Quantification of the average size of giant endosomes in HEK293A WT, HEK-TDP-43-GFP, and HEK-TDP-35-GFP cells. ***, *P* < 0.001 by analysis of variance with Tukey’s *post hoc* test; *N* = 3 (D) HEK-TDP-43-GFP (pseudocolored white) cells were grown in normal conditions before being fixed and immunostained for Rab5, LC3B, or LAMP1 (pseudocolored green) and then successively immunostained for Rab7 (pseudocolored red). *N* = 3; Bar, 10 μm

To further investigate the compositional nature of the giant MVBs in TDP-43-GFP and TDP-35-GFP cells, we performed co-immunostaining of Rab7 along with LC3B, Rab5, or LAMP1 to determine if these were possibly hybrid compartments with autophagosomes, early endosomes, or lysosomes, respectively **(Figure 6D** and **Figure S2B)**. None of these markers strongly colocalized with Rab7 MVB membrane signal, although we did sometimes observe weak colocalization of each marker (most commonly LC3B) with small sections of the MVB membrane **(Figure 6D, Insets)**. Regardless, our giant Rab7-positive vesicular structures appear to predominantly resemble MVBs from a compositional perspective.

### Giant MVBs are proteolytically active compartments

Since our TDP-43-GFP and TDP-35-GFP expressing cells did not show striking GFP signal within giant MVBs, we postulated that these organelles may be proteolytically active and therefore once TDP-43/35-GFP is internalized it is quickly degraded by the resident proteases. Additionally, the lower pH environment of an MVB may impair GFP fluorescence which is sensitive to acidic pH^34^. To address both these issues, we utilized the lysosomotropic agent chloroquine (CHQ), which increases pH within endolysosomal compartments. CHQ could increase detection of any GFP signal within MVBs by increasing GFP fluorescence and impairing resident proteases (as well as fusion with and possible turnover by lysosomes). Indeed, when we treated TDP-43-GFP and TDP-35-GFP cells with CHQ, the GFP signal within giant MVBs became readily apparent **(Figure 7A, Figure S3A)**. While some MVBs exhibited relatively uniform accumulation of TDP-43-GFP **(Figure 7A, row 2)**, we imaged some MVBs in which TDP-43-GFP signal appeared in invagination-like structures around the perimeter of the MVB membrane **(Figure 7A, row 3)**. Additionally, when we transfected TDP-43-GFP and TDP-35-GFP cells with NEDD4-mCherry and then treated the cells with CHQ, we found NEDD4-mCherry colocalized with the TDP-43-GFP and TDP-35-GFP invagination events around the MVB membrane **(Figure S3B)**.

**Figure 7:**
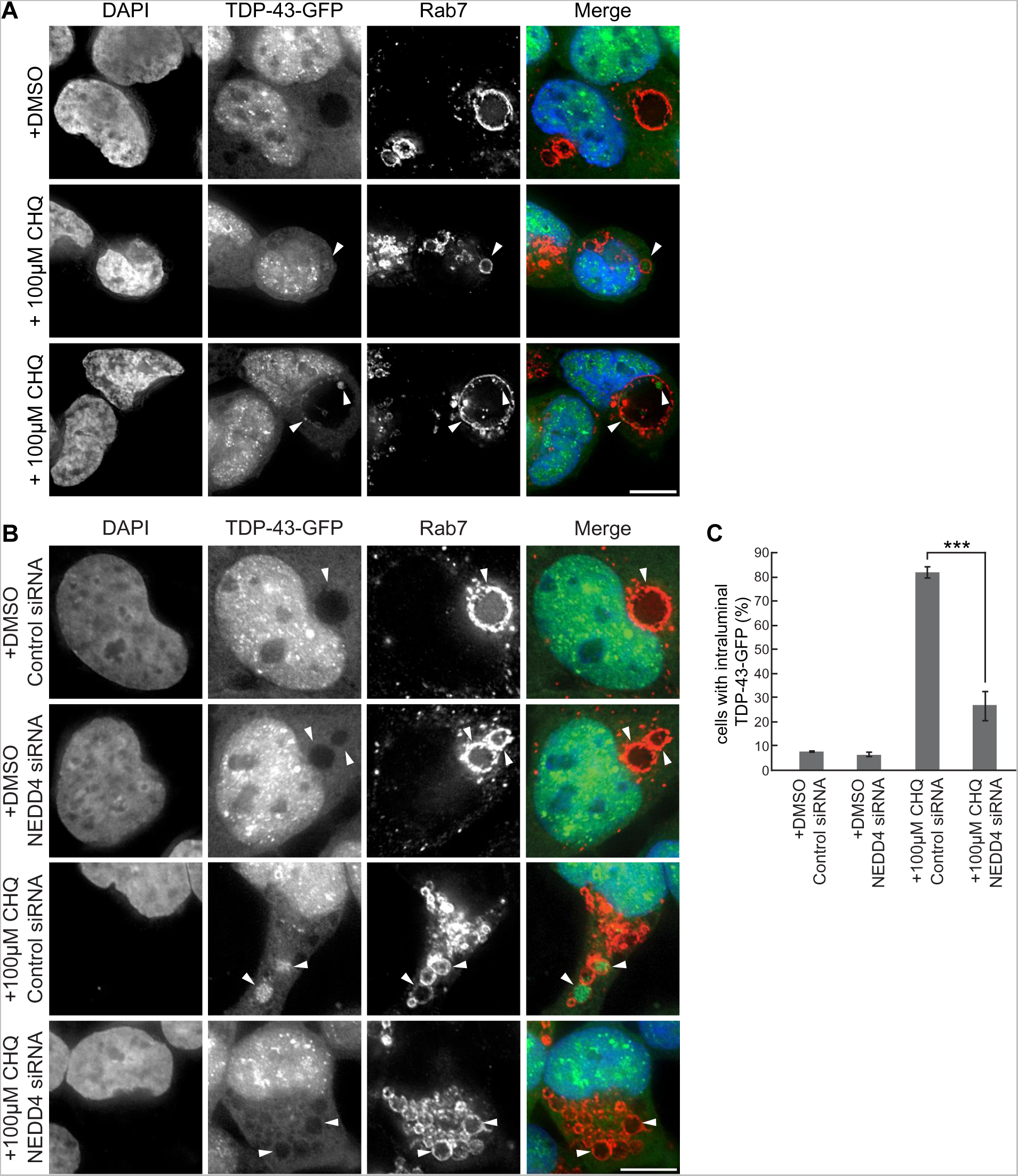
Giant MVBs are proteolytically active compartments (A) HEK-TDP-43-GFP cells were grown in media containing solvent control (DMSO) or 100μM CHQ for 18 hours before being fixed and immunostained for Rab7. *N* = 3; Bar, 10 µm (B) HEK-TDP-43-GFP cells were transfected with siRNAs targeting NEDD4 or non-coding control and grown in normal conditions for 48 hours. Cells were then treated with solvent control (DMSO) or 100μM CHQ for 18 hours before being fixed and immunostained for Rab7. *N* = 3; Bar, 10 µm (C) Quantification of data from (B) showing percentage of cells with enriched TDP-43-GFP signal within Rab7-stained MVBs. ***, *P* < 0.001 by analysis of variance with Tukey’s *post hoc* test; *N* = 2; Bar, 10 µm

We next determined whether NEDD4 knockdown prevented TDP-43 accumulation within giant MVBs. We knocked down NEDD4 in TDP-43-GFP cells and then treated them with CHQ before performing immunofluorescence of Rab7. We found that NEDD4 knockdown significantly decreased the percentage of cells with TDP-43-GFP within MVBs after CHQ treatment **(Figure 7B and 7C)**. This data supports our model that TDP-43 is ubiquitinated by NEDD4 which targets it for internalization into MVBs by the ESCRT complex.

## Discussion

below, we hypothesize that TDP-43, possibly aided by E3-ligase adaptor function (e.g., Rim8 in yeast), is ubiquitinated by Rsp5/NEDD4 which facilitates TDP-43 interaction with various ESCRT complex subunits via their Ub-binding domains. TDP-43 is then incorporated within MVB intraluminal vesicles via ESCRT-mediated invagination. TDP-43 could be degraded either in MVBs^35^, or following subsequent trafficking to vacuoles/lysosomes **(Figure 8)**. Conceptually, this resembles endosomal microautophagy (eMI), which has been described for some cytosolic soluble proteins in human cells, fruit flies, fission yeast (*S. pombe*), but not (at least until now) budding yeast (*S. cerevisiae*)^36–38^. eMI usually involves the Hsc70 chaperone binding a KFERQ-like motif within the substrate protein, which then targets it to the MVB via Hsc70 binding of phosphatidylserine residues on the MVB membrane^39^. Substrates are then internalized into intraluminal vesicles in a manner reliant on components of ESCRT-I, II and III complexes and Vps4^40–42^. Interestingly, under starvation conditions, Hsc70 and ESCRT-I become dispensable for eMI^43^. Notably, knockdown of Tsg101 (ESCRT-I) and Vps24 (ESCRT-III) in HeLa cells results in the accumulation of TDP-43 cytoplasmic aggregates, though this was assumed to reflect macroautophagic defects^44^. TDP-43 also harbors a KFERQ motif and is bound by Hsc70^45^. In *S. pombe*, transport of cytosolic hydrolases to vacuoles occurs via a macroautophagy independent, ESCRT and Ubiquitination-dependent mechanism termed the Nbr1-mediated vacuolar targeting pathway (NVT)^46^. Specifically, Nbr1, three E3 Ub-ligases (two of which are Rsp5 homologs), and a Ub-binding domain within the ESCRT-0 protein Vps27, aid this vacuolar targeting process. Collectively, these studies provide additional support for our model, and highlight the importance of assessing the role of Hsc70, TDP-43’s KFERQ motif and ESCRT Ub-binding functions in future work.

**Figure 8:**
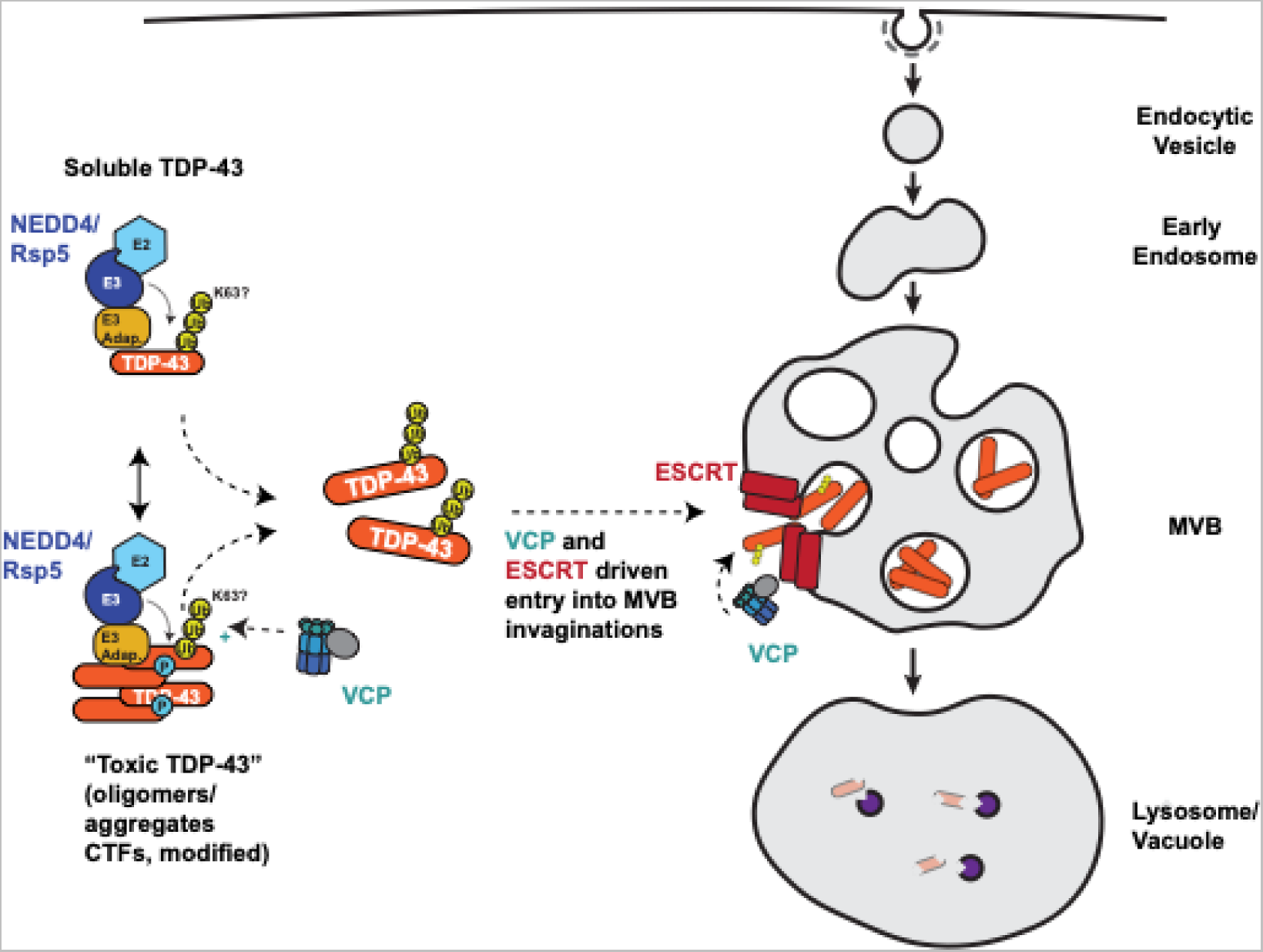
Potential model of Rsp5/NEDD4 and E SCRT-dependent clearance of cytoplasmic TDP-43.

Alternative models of TDP-43 endolysosomal clearance merit consideration. KFERQ in which unfolded proteins are directly translocated across the lysosomal membrane through a LAMP2A translocation pore. While clearance of TDP-43 by CMA has recently been reported^47^, this pathway is absent in yeast, and makes no use of ESCRT components in human cells, thus it is unlikely to relate to the turnover mechanism described in our studies^59, 60^. In contrast, the ESCRT complex, aided by Rsp5 and the Ssh4 adaptor, can also drive vacuole membrane invagination and degradation of vacuolar membrane proteins in yeast^48^. However, *ssh4Δ* yeast cells do not show altered TDP-43 levels (**Figure S1**, despite co-localizing in TDP-43 aggregates (**Figure 3D**). Also, TDP-43 accumulation within Rab7 positive, LAMP1 negative organelles (**Figure 7, Figure S3**) is more consistent with a MVB than lysosomal (vacuolar equivalent) targeting.

Endolysosomal dysfunction may play a key role in the ALS disease mechanism, and possibly contributes to cytoplasmic TDP-43 pathology. Several genes mutated in familial ALS cases (*ALS2*^49, 50^*, FIG4*^51^*, CHMP2B*^52, 53^) have clear endolysosomal functions with *ALS2* being a Rab5 guanine exchange factor (GEF), *FIG4* being involved in PI(3)P synthesis (critical for early endosome, and autophagosome, biogenesis), and *CHMP2B* being an ESCRT-III complex factor. Another familial ALS gene and kinase, *TBK1*^54, 55^, phosphorylates endosomal proteins (including Rab7), may modulate Rab7 GTPase activity^56^, co-localizes with early endosome markers^57^ and leads to severe defects in endolysosomal trafficking and lysosomal acidification when absent in motor neurons^22^. Also, the most common familial ALS mutant gene, *C9ORF72,* harbors a DENN domain which is associated with Guanine exchange factors (GEFs) for Rab GTPases^58, 59^. C9orf72 also colocalizes with early endosome and MVB markers (including in ALS patient motor neurons) and its depletion impairs endolysosomal flux^5947–49^. Finally, partial impairment of *TBK1* activity via an ALS-associated R228H point mutation, combined with (GA)_100_ peptide endolysosomal flux^60^. It is worth noting that TBK1 and C9ORF72 have also been linked to macroautophagic activities^56, 61–63^. Regardless, these studies indicate that endolysosomal dysfunction may lead to cytoplasmic TDP-43 pathology, likely due to impaired TDP-43 clearance.

Simultaneously, a certain cytoplasmic threshold of TDP-43 pathology may also inhibit endolysosomal flux. Overexpression of TDP-43 alleles increasingly prone to cytoplasmic mis-localization and aggregation correlates closely with endocytosis rate defects ^64, 65^. Cytoplasmic TDP-43 aggregation causes dysregulation and lowered abundance of Hsc70-4 at neuromuscular junctions, which drives synaptic vesicle recycling, and endocytosis-dependent process^66^. This could underlie ALS motor neuron defects such as glutamate excitotoxicity, where elevated concentrations of glutamate receptors at synapses are observed^67^. Additionally, Bio-ID proximity labeling data specific to aggregation-prone TDP-43 C-terminal fragments exhibited strong interaction with vesicular transport proteins (most significant of all GO-term categories identified), including endolysosomal and Golgi-ER related pathways^68^. Here, and in our prior work, microscopy confirms that numerous endolysosomal trafficking proteins are sequestered in yeast TDP-43 cytoplasmic aggregates^64^. Thus, while endolysosomal clearance of TDP-43 may be a common response to keep cytoplasmic TDP-43 pathology in check, this pathway might be overwhelmed by accumulation of TDP-43 above a certain cytoplasmic threshold, leading to sequestration of key endolysosomal factors in TDP-43 aggregates. This could lead to a positive feedback loop of further impairment of TDP-43 clearance, and greater sequestration of endolysosomal factors, possibly resulting in compensatory clearance mechanisms being induced (e.g., autophagy) as recently suggested^22^. The appearance of giant Rab7-positive MVB-like structures in our HEK293 TDP-43-GFP and TDP-35-GFP models further supports a defective endolysosomal trafficking process related to TDP-43 pathology. Interestingly, other giant (>2µm) vesicular structures of presumed endolysosomal origin have been observed in ALS relevant contexts. These include *FIG4* mutant mouse fibroblasts, where giant LAMP-2 positive organelles were seen in approximately 40% of cells (versus 5% in WT)^69^. In “wobbler” mutant mice, which develop a progressive ALS-like motor neuron degeneration phenotype due to homozygous Vps54 Q967L mutations (impairs endosome-Golgi trafficking), giant endosomal vesicles were observed in motor neurons, which were Rab7-positive and mostly LC3-negative, mimicking our observations. Amyloid-Precusor Protein (APP; implicated in Alzheimer’s) also co-localized in these structures. These giant endosomal vesicles were also observed in almost half of spinal motor cord neurons from sporadic ALS patients^70^. Finally, in the cortex cells of mice over-expressing a (GA)_100_ peptide, which also exhibited TDP-43 pathology, both EEA1 (early endosome marker) and Rab7 (MVB marker) positive giant endosomes were observed. The frequency of these vesicles was increased further in a *TBK1* hypomorphic background, consistent with the previously discussed endolysosomal trafficking defects of *TBK1* mutants. While we have not examined LAMP2, EEA1 or APP here, these studies all point to a gross defect in endolysosomal trafficking in various ALS models and ALS patient samples. Consistent with the notion that TDP-43 cytoplasmic accumulation may be involved, giant MVBs are observed in >30% of HEK293 cells where endogenous TDP-43 and TDP-43-GFP are co-expressed (2-fold WT levels), whereas WT HEK293 cells display this in <5% of cells (**Figure 6B**). Determining whether simple sequestration of endolysosomal factors within TDP-43 cytoplasmic aggregates, or other TDP-43 mediated inhibitory mechanisms impact endolysosomal trafficking, and in particular giant MVB formation (see^64^ and below) are areas of future interest.

Approximately 600-700 E3 ligases exist in human cells, several of which (Parkin, Cul2, RNF4, Znf179) have been previously linked via candidate-based studies to TDP-43 ubiquitination. Overexpression of both TDP-43 and Parkin in rat brains led to co-localization of both proteins in cortical neurons, though Parkin expression actually increased cytoplasmic TDP-43 mislocalization. Additionally, while Parkin could ubiquitinate TDP-43 *in vitro* with both K48 and K63-linked Ub chains, *in vivo* data did not reveal any impact on TDP-43 ubiquitination nor TDP-43 protein levels^71^. The VHL/Cul2 E3 ligase complex ubiquitinates the 35kDa CTF TDP-43 *in vitro*, co-localizes with TDP-43 inclusions in HEK293 and in ALS patient oligodendrocytes, and facilitates proteasomal-dependent turnover of CTFs *in vivo* based on genetic studies^17^. No impact of VHL/Cul2 on full length TDP-43 levels was observed. RNF4 binds TDP-43 when both proteins are overexpressed. Under heat stress conditions, endogenous TDP-43 exhibits *in vivo* ubiquitination that decreases following RNF4 knockdown, though again no impact on TDP-43 levels was observed. Finally, in mice and N2a cells, Znf179 binds and ubiquitinates TDP-43 *in vivo* and *in vitro* based on genetic data, and aids turnover of endogenous or over-expressed TDP-43 when over-expressed. An increase in cytoplasmic TDP-43 aggregates was also observed in Znf179 KO mice brain tissue^18^. To our knowledge, Rsp5/NEDD4 is the first E3 ligase identified via unbiased methods to impact TDP-43 pathology. Besides demonstrating that Rsp5/NEDD4 regulates TDP-43 levels, stability and aggregation and ubiquitination status, NEDD4 is also the first E3 ligase shown to impact TDP-43 toxicity and targeting to a specific organelle (giant MVBs). Thus, NEDD4 appears to be a consequential regulator of TDP-43 proteostasis, though we cannot rule out that TDP-43 clearance, and other functions, may be regulated by multiple E3 ligases and adaptors.

Like TDP-43, α-synuclein, a Lewy body component present in Parkinson’s, Dementia with Lewy bodies and Multiple System Atrophy, also degrades in yeast and human cell culture models via an autophagy and proteasomal independent mechanism. This too relies on ESCRT and Rsp5/NEDD4 function, with NEDD4-mediated K63-ubiquitination of α-synuclein having been demonstrated^72, 73^. Fly and rat models of Parkinson’s, featuring ectopic α-synuclein expression, show suppression and enhancement of α-synuclein pathological effects with Nedd4 over-expression or knockdown respectively^74^. We speculate that many proteins besides TDP-43 and α-synuclein are likely cleared via an Rsp5/NEDD4 and ESCRT-regulated endolysosomal pathway.

Distinct TDP-43 protein isoforms arising from ALS-associated mutations, alternative splicing or post-translational cleavage exhibit divergent protein half-lives from WT TDP-43^757677–80^, implying distinct clearance pathways are likely used. Thus, TDP-43 isoform, misfolding, aggregation and modification status are likely determining factors for utilization of the endolysosomal clearance pathway. This could be regulated in part by VCP/p97, a Ub-segregase and AAA-ATPase chaperone, which also facilitates TDP-43 clearance^65^, and which can facilitate entry of ubiquitinated substrates into MVBs^81–84^. Addressing how distinct isoforms of TDP-43 are degraded is an important area of future research as it may allow selective turnover of critical disease-associated isoforms while preserving normal TDP-43 and its associated cellular functions.

## Materials and Methods

### Yeast strains and growth conditions

Untransformed yeast strains were cultured at 30°C with yeast extract-peptone-dextrose (YPD) medium. Strains transformed with plasmids were grown in synthetic defined (SD) medium with appropriate nutrients for plasmid selection. In liquid culture, strains expressing *GAL1*-driven TDP-43 from a plasmid (or vector control) were initially cultured overnight in 2% sucrose and then back diluted to an optical density at 600 nm (OD_600_) of 0.1-0.15 in 0.25% galactose and 1.75% sucrose medium, followed by growth at 30°C to mid-logarithmic growth phase (OD_600_, 0.4 to 0.7). Transformations were performed by a standard lithium acetate method. All yeast strain and plasmids utilized in the study are listed in **Supplemental Table 3**.

### Dot blot genetic screen

To identify genes that affect TDP-43 protein levels, we developed a yeast genetic screening method utilizing high throughput yeast colony dot blots. The yeast gene deletion library was transformed via a high-throughput transformation approach ^81^ with a single-copy plasmid encoding TDP-43-YFP under the control of a galactose regulatable promoter. An outgrowth of transformants was then performed in 96-well suspension culture plates for 4 days following transformation in SD media lacking uracil and supplemented with 2% glucose. These yeast strains were then pinned in 384-well format onto agar plates containing the appropriate selection with 2% sucrose as the carbon source using a Singer ROTOR HDA. After growing for 48 hours at 30°C, the yeast colonies were then re-pinned in 384-well format onto agar plates that had been overlaid with nitrocellulose. These nitrocellulose-coated agar plates contained appropriate plasmid selection and 1.75% sucrose and 0.25% galactose to induce the expression of TDP-43-YFP. After growth for 8 hours at 30°C, the nitrocellulose was removed from the agar plate and placed onto filter paper that had been soaked in lysis buffer (350mM NaOH, 0.1% SDS, and 1% 2-mercaptoethanol) for 15 minutes to lyse the cells *in situ*. Blots were then washed three times with 1x TBS+0.1% Tween-20 to remove cell debris. Blots were then blocked using 5% non-fat dry milk (NFDM) for 45 minutes at room temperature and incubated with primary antibodies: anti-GFP (catalog number ab290; Abcam; dilution 1:5000) and anti-GAPDH (catalog number MA5-15738; Invitrogen; dilution 1:5000) for 16 hours at 4°C. Blots were then washed three times with 1x TBS+0.1% Tween-20 for 5 minutes each. Blots were incubated with secondary antibodies: IRDye 800CW goat anti-rabbit (catalog number 925-32211; LI-COR; dilution 1:20000) and IRDye 680RD donkey anti-mouse (catalog number 926-68072; LI-COR; dilution 1:20000) at room temperature for 1 hour. Blots were then washed four times with 1x TBS+0.1% Tween-20 for 5 minutes each. Blots were then kept in 1x PBS until being imaged using a LI-COR Odyssey infrared imaging system.

### Western blotting

Western blotting was conducted as described previously^20^. Primary antibodies were as follows: anti-GFP (catalog number: QAB10298; enQuirebio; dilution 1:5000), anti-GAPDH (anti-glyceraldehyde-3-phosphate dehydrogenase; catalog number MA5-15738; Invitrogen; dilution 1:5000), anti-TDP-43 (catalog number 10782-2-AP; Proteintech; dilution 1:5000), anti-NEDD4 (catalog number 21698-1-AP; Proteintech; dilution 1:1000), anti-Ubiquitin (catalog number sc-8017; Santa Cruz; dilution 1:1000), and anti-mCherry (catalog number ab167453, Abcam; dilution 1:1000). The results of the quantification of key Western blotting datasets from throughout the paper are listed in **Supplementary Table 2**.

### Cycloheximide chase assays

For TDP-43 protein stability assays in yeast, TDP-43 expression was halted by the addition of cycloheximide at 0.2 mg/ml to the growth media. Protein lysates were obtained via a standard NaOH method and examined via Western blot. For TDP-43-GFP protein stability assays in HEK293A cells, TDP-43-GFP expression was halted by the addition of cycloheximide at 0.3 mg/ml to the growth media. Protein lysates were generated by resuspending cell pellets in a standard SDS loading buffer followed by incubation at 95°C for 10 minutes and examined via Western blot.

### Yeast fluorescence microscopy

The methods have been described previously^20^. Briefly, mid-log live yeast cells (OD_600_ 0.4–0.7) were rapidly examined at room temperature on glass slides (Globe Scientific, cat:1301) with 1.5 coverslips (VWR, cat: 48366-227) using a DeltaVision Elite microscope with a ×100 oil-immersion (1.515; Cargille, cat: 16245) objective (NA 1.40), and 15-bit PCO Edge sCMOS camera. Z-stack data (10 slices, 0.4μm each) was collected. Images were subject to deconvolution using standard parameters (Enhanced ratio, 10 cycles, medium filtering) using Softworx 7.0.0 (Deltavision software), followed by maximum intensity Z-stack projection. All image analysis was done using Fiji software^82^. TDP-43 foci co-localization with different proteins and TDP-43 foci per cell was scored manually, in a blind manner, in a minimum of 50 cells (in which each cell may contain multiple foci). Cell and foci counts for all microscopy data (yeast and human) are listed in **Supplementary Table 2**. All shown images are representative of >3 biological replicates, in which each biological replicate involves analysis of a minimum of 50 cells.

### Immunoprecipitation analyses

For immunoprecipitation analyses in yeast cells, 10 ODs of mid-logarithmic growth phase yeast were pelleted by centrifugation at 13,000 x *g* at 4°C and resuspended in 1 ml of radioimmunoprecipitation assay cell lysis buffer. 50% volume of disruption beads was added, and the cells were vortexed for 2 min before being incubated on ice for 3 min. This process was repeated twice. The cells were centrifuged at 17,000 x *g* for 10 min at 4°C to separate unlysed cells and insoluble cell debris. Supernatants were applied to GFP-Trap magnetic agarose beads (catalog number: gtma, ChromoTek) and nutated for 2 hours at 4°C. Beads were then washed 4 times with wash buffer before being resuspended in standard SDS loading buffer, boiled at 95°C for 10 minutes, and examined via Western blot. For immunoprecipitation analysis in HEK293A cells, 10cm dishes of cells stably expressing TDP-43-GFP were transfected with NEDD4-mCherry or mock control and grown to 80-90% confluence over 2 days. Cells were collected via centrifugation and washed in cold 1x DPBS twice before being resuspended in radioimmunoprecipitation assay cell lysis buffer. The cells were centrifuged at 17,000 x *g* for 10mins at 4°C to separate insoluble cell debris. Supernatants were applied to RFP-Trap magnetic agarose beads (catalog number: rtma, ChromoTek) and nutated for 2 hours at 4°C. Beads were then washed 4 times with wash buffer before being resuspended in standard SDS loading buffer, boiled at 95°C for 10 minutes, and examined via Western blot

### Yeast serial dilution growth assays

Yeast strains transformed with plasmids expressing TDP-43 or an empty vector were grown overnight as described above, back diluted to an OD_600_ of 0.2, serially diluted (1:5 dilutions), and spotted onto identical agar media. All images are representative of those from a minimum of three biological replicates imaged on the same day.

### Culture of HEK293A cells and HEK293A cells expressing TDP-43–GFP or TDP-35–GFP

TDP-43–GFP and a 35kDa C-terminal TDP-43 fragment (TDP-35–GFP; amino acids 90 to 414, a cytoplasmic aggregation-prone allele mimicking TDP-43 cleavage by caspases^83^) were stably integrated by retroviral means (pQCXIH vector, Hygromycin selection) were stably integrated into HEK293A ells by retroviral means as previously described^19^. Resulting expression of TDP-43-GFP and TDP-35-GFP are comparable to the level of endogenous TDP-43, meaning both lines have about double the amount of TDP-43 species (endogenous and GFP-tagged versions) when compared to HEK293A WT cells. HEK293A WT, HEK-TDP-43-GFP, and HEK-TDP-35-GFP cells were cultured at 37°C in Dulbecco modified Eagle medium (catalog number 10-013-CV; Corning) with 10% fetal bovine serum (catalog number 26140079; Gibco), and 50 U/ml penicillin and 50 μg/ml streptomycin (catalog number 15140-122; Gibco). Plasmid transfections were performed using Lipofectamine 3000 according to the manufacturer’s protocol (catalog number L3000008; Thermo Scientific). siRNA transfections were performed using Lipofectamine RNAiMAX according to the manufacturer’s protocol (catalog number 13778100; Thermo Scientific).

### Cell line viability assessment

HEK293A, HEK-TDP-43-GFP and HEK-TDP-35-GFP cells were seeded in 6-well plates at 1×10^5^ cells per well. Cells were then transfected with 2.5µg NEDD4-mCherry or VPS4^E228Q^ plasmid, or 25 pmol NEDD4 siRNA, or appropriate controls and incubated for 48 hours. Cell numbers were manually counted using a hemocytometer and normalized to the WT control wells.

### Ubiquitination assay via immunoprecipitation

To assess TDP-43-GFP ubiquitination in HEK-TDP-43-GFP cells, 10cm dishes of cells were transfected with NEDD4-mCherry, siRNAs targeting NEDD4, or mock control and grown to 80-90% confluence over 3 days. Cells were collected via centrifugation and washed in cold 1x DPBS twice before being resuspended in urea lysis buffer (8M urea, 50mM Tris-HCl pH 8.0, 75mM NaCl, 100mM N-ethylmaleimide, 1mM β-glycerophosphate, 1mM NaF, 1mM NaV, and EDTA-free protease inhibitor cocktail (Roche Diagnostics). Lysates were applied to GFP-Trap magnetic agarose beads and nutated for 2 hours at 4°C. Beads were then washed 4 times with urea lysis buffer before being resuspended in standard SDS loading buffer, boiled at 95°C for 10 minutes, and examined via Western blot.

### Cell line immunofluorescence

Cells were cultured for 24 h in 8-well chamber slides (catalog number 80821; Ibidi), fixed with 4% paraformaldehyde for 15 minutes, and permeabilized with 0.1% Triton X-100 for 15 min. Cells were incubated with indicated primary antibodies overnight at 4°C after blocking with 5% bovine serum albumin. Cells were then incubated with Alexa fluor conjugated secondary antibody (Thermo Fisher Scientific, 1:1000 dilution) for 1 h at room temperature. After three washes with PBS, cells were stained with 0.5 µg/ml DAPI (catalog number P36931; Thermo Fisher Scientific) and imaged using a Deltavision Elite microscope. Primary antibodies used for the immunofluorescence were: Rab7 (ab137029; Abcam, dilution: 1:200), Rab5 (ab109534; Abcam, dilution: 1:200), LC3B (3868S; Cell Signaling, dilution 1:500), LAMP1 (21997-1-AP; Proteintech, dilution 1:200), and TDP-43 (catalog number 10782-2-AP; Proteintech; dilution 1:200). All shown images are representative of 3 biological replicates, except for Figure 7B which images shown are representative of 2 biological replicates.

## Acknowledgements

This work was funded by grants from the NIH (NIGMS R01GM114564 and NINDS R56NS128110). A.B. received support from the NSF Graduate Research Fellowship Program (DGE-1746060), L.M received support from an NIH training grant (T32GM136536) and S.M received support from the Undergraduate Biology Research Program (UBRP) at the University of Arizona. We thank Matt Kaplan and the Function Genomics Core at the University of Arizona for their help with our genetic screen and Buchan lab members for review and comments on the manuscript.

## Author contributions

A.B. and J.R.B. conceived of the project and wrote the manuscript. L.M. and S.M. aided Rsp5 adaptor experiments. V.A. performed Rsp5 over-expression analyses in yeast. All other experimental data was collected and analyzed by A.B.

---

**Figure S1:**
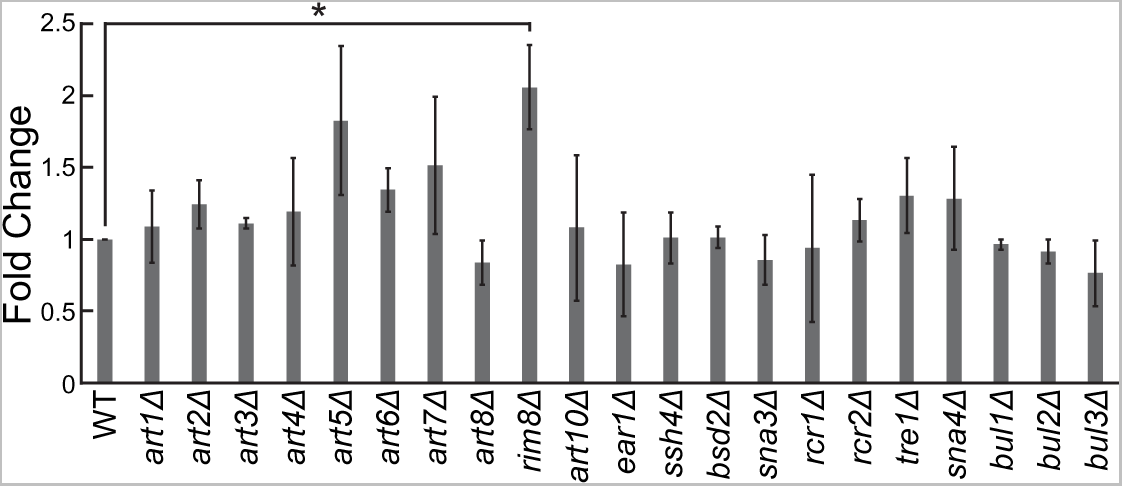
Knockout of the Rsp5 adaptor Rim8 increases TDP-43-YFP levels. Indicated strains were transformed with TDP-43-YFP plasmid and grown to mid-logarithmic growth phase. TDP-43-YFP levels were detected by western blot using anti-TDP-43 antibody and normalized to GAPDH loading control; quantification shown here. *, P < 0.05; ***, *P* < 0.001 analysis of variance with Tukey’s *post hoc* test; *N* = 3, except for strains *art2-4Δ, art8Δ, art10Δ, bsd2Δ, sna3Δ, bul1-2Δ* (N = 2).

**Figure S2:**
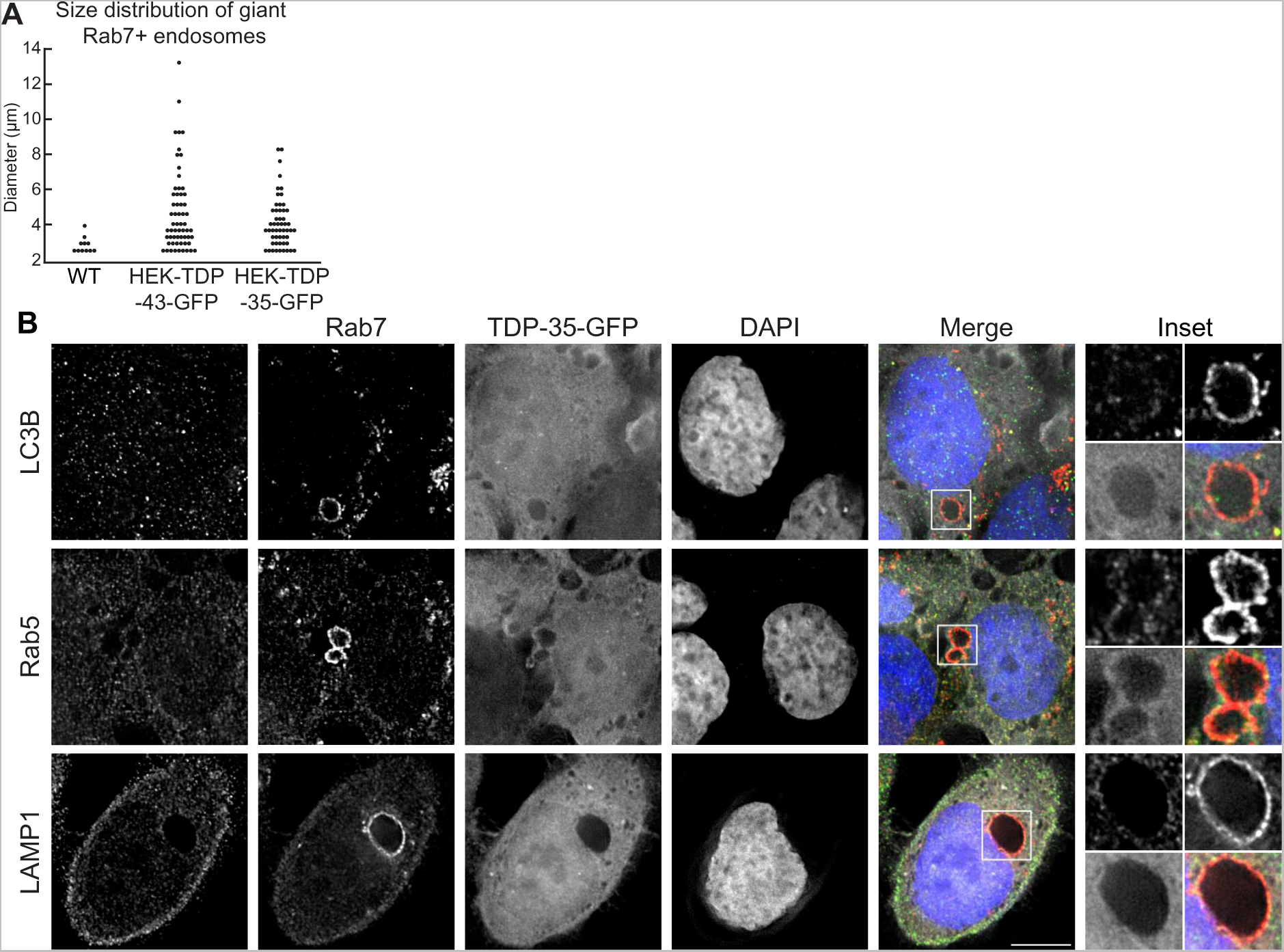
HEK-TDP-43-GFP and HEK-TDP-35-GFP cell lines develop giant MVBs (A) Distribution of giant MVB diameters from WT, HEK-TDP-43-GFP, and HEK-TDP-35-GFP cell lines. Quantification was determined by manually measuring diameter of giant MVBs using Fiji software. *N* = 3 (B) HEK-TDP-35-GFP (pseudocolored white) cells were grown in normal conditions before being fixed and immunostained for Rab5, LC3B, or LAMP1 (pseudocolored green) and then successively immunostained for Rab7 (pseudocolored red). *N* = 3; Bar, 10 µm

**Figure S3:**
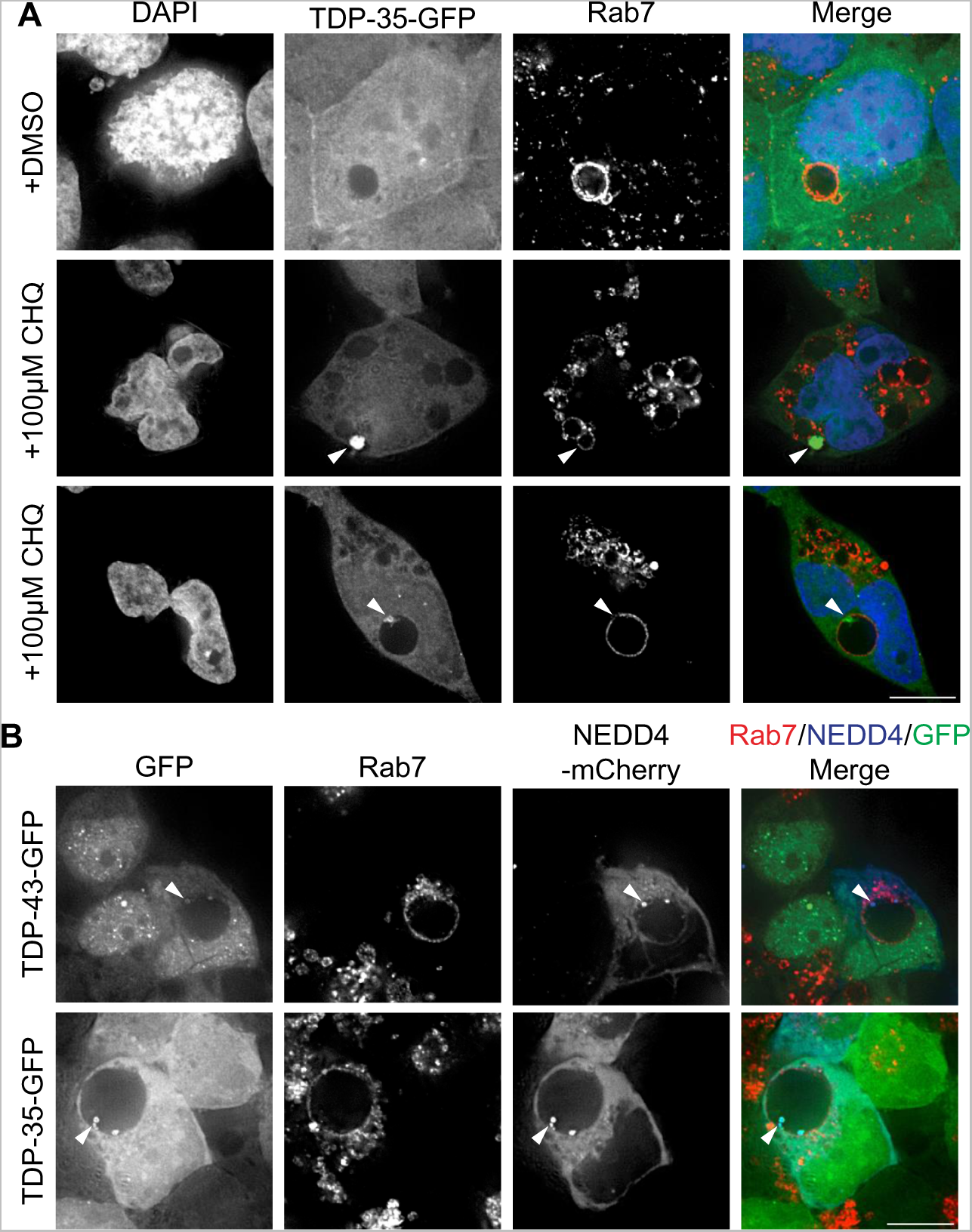
Giant MVBs are proteolytically active compartments (A) HEK-TDP-35-GFP cells were grown in media containing solvent control (DMSO; top row) or 100μM CHQ for 18 hours (next 2 rows, to show phenotypic diversity) before being fixed and immunostained for Rab7. Arrowheads indicate TDP-35 within Rab7-positive giant MVBs. *N* = 3; Bar, 10 µm (B) HEK-TDP-43-GFP and HEK-TDP-35-GFP cells were transfected with NEDD4-mCherry plasmid and grown in normal conditions for 24 hours. Cells were then treated with 100µM CHQ and incubated for 18 hours before being fixed and immunostained for Rab7. Arrowheads indicate TDP-45 and TDP-35-GFP signal co-localized with NEDD4-mCh within Rab7-positive giant MVBs *N* = 3; Bar, 10 µm

## References

1. Taylor, J. P., Brown, R. H. & Cleveland, D. W. Decoding ALS: From genes to mechanism. Nature vol. 539 197–206 Preprint at https://doi.org/10.1038/nature20413 (2016).

2. Ling, S.-C., Polymenidou, M. & Cleveland, D. W. Converging Mechanisms in ALS and FTD: Disrupted RNA and Protein Homeostasis. Neuron 79, 416–438 (2013).

3. Cascella, R. et al. Quantification of the Relative Contributions of Loss-of-function and Gain-of-function Mechanisms in TAR DNA-binding Protein 43 (TDP-43) Proteinopathies. Journal of Biological Chemistry 291, 19437–19448 (2016).

4. Fecto, F. et al. SQSTM1 Mutations in Familial and Sporadic Amyotrophic Lateral Sclerosis. Arch. Neurol. 68, 1440–1446 (2011).

5. Watts, G. D. J. et al. Inclusion body myopathy associated with Paget disease of bone and frontotemporal dementia is caused by mutant valosin-containing protein. Nat Genet 36, 377–381 (2004).

6. Deng, H. X. et al. Mutations in UBQLN2 cause dominant X-linked juvenile and adult-onset ALS and ALS/dementia. Nature 477, 211–215 (2011).

7. Maruyama, H. et al. Mutations of optineurin in amyotrophic lateral sclerosis. Nature 465, 223–226 (2010).

8. Johnson, B. S., McCaffery, J. M., Lindquist, S. & Gitler, A. D. A yeast TDP-43 proteinopathy model: Exploring the molecular determinants of TDP-43 aggregation and cellular toxicity. Proc Natl Acad Sci U S A 105, 6439–6444 (2008).

9. Li, Y. et al. A Drosophila model for TDP-43 proteinopathy. Proc Natl Acad Sci U S A 107, 3169–3174 (2010).

10. Bilican, B. et al. Mutant induced pluripotent stem cell lines recapitulate aspects of TDP-43 proteinopathies and reveal cell-specific vulnerability. Proceedings of the National Academy of Sciences 109, 5803–5808 (2012).

11. Wang, I.-F. et al. Autophagy activators rescue and alleviate pathogenesis of a mouse model with proteinopathies of the TAR DNA-binding protein 43. Proceedings of the National Academy of Sciences 109, 15024–15029 (2012).

12. SJ, B. et al. Amelioration of toxicity in neuronal models of amyotrophic lateral sclerosis by hUPF1. Proc Natl Acad Sci U S A 112, 7821–7826 (2015).

13. Becker, L. A. et al. Therapeutic reduction of ataxin-2 extends lifespan and reduces pathology in TDP-43 mice. Nature 2017 544:7650 544, 367–371 (2017).

14. Castillo, K. et al. Trehalose delays the progression of amyotrophic lateral sclerosis by enhancing autophagy in motoneurons. Autophagy 9, 1308–1320 (2013).

15. Hans, F. et al. UBE2E ubiquitin-conjugating enzymes and ubiquitin isopeptidase y regulate TDP-43 protein ubiquitination. Journal of Biological Chemistry 289, 19164–19179 (2014).

16. Hebron, M. L. et al. Parkin ubiquitinates tar-DNA binding protein-43 (TDP-43) and promotes its cytosolic accumulation via interaction with histone deacetylase 6 (HDAC6). Journal of Biological Chemistry 288, 4103–4115 (2013).

17. Uchida, T. et al. CUL2-mediated clearance of misfolded TDP-43 is paradoxically affected by VHL in oligodendrocytes in ALS. Sci Rep 6, 1–19 (2016).

18. Lee, Y. C. et al. Znf179 E3 ligase-mediated TDP-43 polyubiquitination is involved in TDP-43-ubiquitinated inclusions (UBI) (+)-related neurodegenerative pathology 11 Medical and Health Sciences. J Biomed Sci 25, (2018).

19. Liu, G. et al. Endocytosis regulates TDP-43 toxicity and turnover. doi:10.1038/s41467-017-02017-x.

20. Liu, G. et al. Cdc48/VCP and Endocytosis Regulate TDP-43 and FUS Toxicity and Turnover. Mol Cell Biol 40, 1–18 (2020).

21. Leibiger, C. et al. TDP-43 controls lysosomal pathways thereby determining its own clearance and cytotoxicity. Hum Mol Genet 27, 1593–1607 (2018).

22. Hao, J. et al. Loss of TBK1 activity leads to TDP-43 proteinopathy through lysosomal dysfunction in human motor neurons. bioRxiv 2021.10.11.464011 (2021) doi:10.1101/2021.10.11.464011.

23. Zacchi, L. F. et al. Early-onset torsion dystonia: a novel high-throughput yeast genetic screen for factors modifying protein levels of torsinAΔE. Dis Model Mech 10, 1129 (2017).

24. Fang, N. N. et al. Rsp5/Nedd4 is the main ubiquitin ligase that targets cytosolic misfolded proteins following heat stress. Nature Cell Biology 2014 16:12 16, 1227–1237 (2014).

25. Herrador, A., Herranz, S., Lara, D. & Vincent, O. Recruitment of the ESCRT Machinery to a Putative Seven-Transmembrane-Domain Receptor Is Mediated by an Arrestin-Related Protein. Mol Cell Biol 30, 897–907 (2010).

26. Wen, R., Li, J., Xu, X., Cui, Z. & Xiao, W. Zebrafish Mms2 promotes K63-linked polyubiquitination and is involved in p53-mediated DNA-damage response. DNA Repair (Amst) 11, 157–166 (2012).

27. Simões, V. et al. Redox-sensitive E2 Rad6 controls cellular response to oxidative stress via K63-linked ubiquitination of ribosomes. Cell Rep 39, 110860 (2022).

28. Katzmann, D. J., Babst, M. & Emr, S. D. Ubiquitin-Dependent Sorting into the Multivesicular Body Pathway Requires the Function of a Conserved Endosomal Protein Sorting Complex, ESCRT-I. Cell 106, 145–155 (2001).

29. Rothfels, K. et al. Components of the ESCRT Pathway, DFG16, and YGR122w Are Required for Rim101 to Act as a Corepressor with Nrg1 at the Negative Regulatory Element of the DIT1 Gene of Saccharomyces cerevisiae. Mol Cell Biol 25, 6772–6788 (2005).

30. Sardana, R. & Emr, S. D. Membrane Protein Quality Control Mechanisms in the Endo-Lysosome System. Trends Cell Biol 31, 269–283 (2021).

31. Neumann, S. et al. Formation and nuclear export of tRNA, rRNA and mRNA is regulated by the ubiquitin ligase Rsp5p. EMBO Rep 4, 1156–1162 (2003).

32. Votteler, J. et al. Designed proteins induce the formation of nanocage-containing extracellular vesicles. Nature 540, 292–295 (2016).

33. Von Bartheld, C. S. & Altick, A. L. Multivesicular Bodies in Neurons: Distribution, Protein Content, and Trafficking Functions. Prog Neurobiol 93, 313 (2011).

34. Shinoda, H., Shannon, M. & Nagai, T. Fluorescent Proteins for Investigating Biological Events in Acidic Environments. Int J Mol Sci 19, (2018).

35. Krause, G. J. & Cuervo, A. M. Assessment of mammalian endosomal microautophagy. Methods Cell Biol 164, 167–185 (2021).

36. Sahu, R. et al. Microautophagy of Cytosolic Proteins by Late Endosomes. Dev Cell 20, 131–139 (2011).

37. Mukherjee, A., Patel, B., Koga, H., Cuervo, A. M. & Jenny, A. Selective endosomal microautophagy is starvation-inducible in Drosophila. Autophagy 12, 1984–1999 (2016).

38. Liu, X.-M. et al. ESCRTs Cooperate with a Selective Autophagy Receptor to Mediate Vacuolar Targeting of Soluble Cargos. Mol Cell 59, 1035–1042 (2015).

39. Morozova, K. et al. Structural and Biological Interaction of hsc-70 Protein with Phosphatidylserine in Endosomal Microautophagy. J Biol Chem 291, 18096–18106 (2016).

40. Tekirdag, K. & Cuervo, A. M. Chaperone-mediated autophagy and endosomal microautophagy: Joint by a chaperone. Journal of Biological Chemistry vol. 293 5414–5424 Preprint at https://doi.org/10.1074/jbc.R117.818237 (2018).

41. Mukherjee, A., Patel, B., Koga, H., Cuervo, A. M. & Jenny, A. Selective endosomal microautophagy is starvation-inducible in Drosophila. Autophagy 12, 1984–1999 (2016).

42. Sahu, R. et al. Microautophagy of Cytosolic Proteins by Late Endosomes. Dev Cell 20, 131–139 (2011).

43. Mejlvang, J. et al. Starvation induces rapid degradation of selective autophagy receptors by endosomal microautophagy. J Cell Biol 217, 3640–3655 (2018).

44. Filimonenko, M. et al. Functional multivesicular bodies are required for autophagic clearance of protein aggregates associated with neurodegenerative disease. Journal of Cell Biology 179, 485–500 (2007).

45. Huang, C. C. et al. Metabolism and mis-metabolism of the neuropathological signature protein TDP-43. J Cell Sci 127, 3024–3038 (2014).

46. Liu, X.-M. et al. ESCRTs Cooperate with a Selective Autophagy Receptor to Mediate Vacuolar Targeting of Soluble Cargos. Mol Cell 59, 1035–1042 (2015).

47. Ormeño, F. et al. Chaperone Mediated Autophagy Degrades TDP-43 Protein and Is Affected by TDP-43 Aggregation. Front Mol Neurosci 13, 19 (2020).

48. Zhu, L., Jorgensen, J. R., Li, M., Chuang, Y. S. & Emr, S. D. ESCRTS function directly on the lysosome membrane to downregulate ubiquitinated lysosomal membrane proteins. Elife 6, (2017).

49. Hadano, S. et al. A gene encoding a putative GTPase regulator is mutated in familial amyotrophic lateral sclerosis 2. Nat Genet 29, 166–173 (2001).

50. Kunita, R. et al. Homo-oligomerization of ALS2 through its unique carboxyl-terminal regions is essential for the ALS2-associated Rab5 guanine nucleotide exchange activity and its regulatory function on endosome trafficking. J Biol Chem 279, 38626–38635 (2004).

51. Chow, C. Y. et al. Deleterious Variants of FIG4, a Phosphoinositide Phosphatase, in Patients with ALS. doi:10.1016/j.ajhg.2008.12.010.

52. Parkinson, N. et al. ALS phenotypes with mutations in CHMP2B (charged multivesicular body protein 2B). Neurology 67, 1074–1077 (2006).

53. Skibinski, G. et al. Mutations in the endosomal ESCRTIII-complex subunit CHMP2B in frontotemporal dementia. Nat Genet 37, 806–8 (2005).

54. Cirulli, E. T. et al. Exome sequencing in amyotrophic lateral sclerosis identifies risk genes and pathways. Science 347, 1436–1441 (2015).

55. Freischmidt, A. et al. Haploinsufficiency of TBK1 causes familial ALS and fronto-temporal dementia. Nat Neurosci 18, 631–636 (2015).

56. Heo, J. M. et al. RAB7A phosphorylation by TBK1 promotes mitophagy via the PINK-PARKIN pathway. Sci Adv 4, (2018).

57. Chau, T.-L. et al. A role for APPL1 in TLR3/4-dependent TBK1 and IKKε activation in macrophages. J Immunol 194, 3970–3983 (2015).

58. Levine, T. P., Daniels, R. D., Gatta, A. T., Wong, L. H. & Hayes, M. J. The product of C9orf72, a gene strongly implicated in neurodegeneration, is structurally related to DENN Rab-GEFs. Bioinformatics 29, 499 (2013).

59. Farg, M. A. et al. C9ORF72, implicated in amyotrophic lateral sclerosis and frontotemporal dementia, regulates endosomal trafficking. Hum Mol Genet 23, 3579–3595 (2014).

60. Shao, W. et al. Two FTD-ALS genes converge on the endosomal pathway to induce TDP-43 pathology and degeneration. Science 378, 94–99 (2022).

61. Pilli, M. et al. TBK-1 promotes autophagy-mediated antimicrobial defense by controlling autophagosome maturation. Immunity 37, 223–234 (2012).

62. Webster, C. P. et al. The C9orf72 protein interacts with Rab1a and the ULK1 complex to regulate initiation of autophagy. EMBO J 35, 1656–76 (2016).

63. Nassif, M., Woehlbier, U. & Manque, P. A. The Enigmatic Role of C9ORF72 in Autophagy. Front Neurosci 11, 442 (2017).

64. Liu, G. et al. Endocytosis regulates TDP-43 toxicity and turnover. Nat Commun 8, 2092 (2017).

65. Liu, G. et al. Cdc48/VCP and endocytosis regulate TDP-43 and FUS toxicity and turnover. Mol Cell Biol 40, (2020).

66. Coyne, A. N. et al. Post-transcriptional Inhibition of Hsc70-4/HSPA8 Expression Leads to Synaptic Vesicle Cycling Defects in Multiple Models of ALS. Cell Rep 21, 110–125 (2017).

67. Shi, Y. et al. Haploinsufficiency leads to neurodegeneration in C9ORF72 ALS/FTD human induced motor neurons. Nat Med 24, 313 (2018).

68. Chou, C. C. et al. TDP-43 pathology disrupts nuclear pore complexes and nucleocytoplasmic transport in ALS/FTD. Nat Neurosci 21, 228–239 (2018).

69. Chow, C. Y. et al. Mutation of FIG4 causes neurodegeneration in the pale tremor mouse and patients with CMT4J. Nature 448, 68–72 (2007).

70. Palmisano, R. et al. Endosomal accumulation of APP in wobbler motor neurons reflects impaired vesicle trafficking: implications for human motor neuron disease. BMC Neurosci 12, (2011).

71. Hebron, M. L. et al. Parkin ubiquitinates tar-DNA binding protein-43 (TDP-43) and promotes its cytosolic accumulation via interaction with histone deacetylase 6 (HDAC6). Journal of Biological Chemistry 288, 4103–4115 (2013).

72. Tofaris, G. K. et al. Ubiquitin ligase Nedd4 promotes α-synuclein degradation by the endosomal-lysosomal pathway. Proc Natl Acad Sci U S A 108, 17004–17009 (2011).

73. Tardiff, D. F. et al. Yeast Reveal a “Druggable” Rsp5/Nedd4 Network that Ameliorates-Synuclein Toxicity in Neurons. Science (1979) 342, 979–983 (2013).

74. Davies, S. E. et al. Enhanced ubiquitin-dependent degradation by Nedd4 protects against α-synuclein accumulation and toxicity in animal models of Parkinson’s disease. Neurobiol Dis 64, 79–87 (2014).

75. Flores, B. N. et al. An Intramolecular Salt Bridge Linking TDP43 RNA Binding, Protein Stability, and TDP43-Dependent Neurodegeneration. Cell Rep 27, 1133–1150.e8 (2019).

76. Watanabe, S., Kaneko, K. & Yamanaka, K. Accelerated disease onset with stabilized familial amyotrophic lateral sclerosis (ALS)-linked mutant TDP-43 proteins. Journal of Biological Chemistry 288, 3641–3654 (2013).

77. Pesiridis, G. S., Tripathy, K., Tanik, S., Trojanowski, J. Q. & Lee, V. M. Y. A ‘two-hit’ hypothesis for inclusion formation by carboxyl-terminal fragments of TDP-43 protein linked to RNA depletion and impaired microtubule-dependent transport. J Biol Chem 286, 18845–18855 (2011).

78. Scotter, E. L. et al. Differential roles of the ubiquitin proteasome system and autophagy in the clearance of soluble and aggregated TDP-43 species. J Cell Sci 127, 1263–78 (2014).

79. Brower, C. S., Piatkov, K. I. & Varshavsky, A. Neurodegeneration-associated protein fragments as short-lived substrates of the N-end rule pathway. Mol Cell 50, 161–171 (2013).

80. Berning, B. A. & Walker, A. K. The pathobiology of TDP-43 C-terminal fragments in ALS and FTLD. Front Neurosci 13, 335 (2019).

81. Kirchner, P., Bug, M. & Meyer, H. Ubiquitination of the N-terminal region of caveolin-1 regulates endosomal sorting by the VCP/p97 AAA-ATPase. J Biol Chem 288, 7363–7372 (2013).

82. Jang, H. I., Jang, E. R., Wilson, P. G., Anderson, D. & Galperin, E. VCP/p97 controls signals of the ERK1/2 pathway transmitted via the Shoc2 scaffolding complex: Novel insights into IBMPFD pathology. Mol Biol Cell 30, 1655–1663 (2019).

83. Ritz, D. et al. Endolysosomal sorting of ubiquitylated caveolin-1 is regulated by VCP and UBXD1 and impaired by VCP disease mutations. Nat Cell Biol 13, 1116–1124 (2011).

84. Burana, D. et al. The Ankrd13 Family of Ubiquitin-interacting Motif-bearing Proteins Regulates Valosin-containing Protein/p97 Protein-mediated Lysosomal Trafficking of Caveolin 1. J Biol Chem 291, 6218–6231 (2016).

85. Gietz, R. D. & Schiestl, R. H. Microtiter plate transformation using the LiAc/SS carrier DNA/PEG method. Nature Protocols 2007 2:1 2, 5–8 (2007).

86. Schindelin, J., et al. Fiji: an open-source platform for biological-image analysis. Nature Methods 2012 9:7 **9**, 676–682 (2012).

87. Zhang, Y. J. et al. Aberrant cleavage of TDP-43 enhances aggregation and cellular toxicity. Proc Natl Acad Sci U S A 106, 7607–7612 (2009).

